# Pre-existing immunity to influenza viruses through infection and/or vaccination leads to viral mutational signatures associated with unique immune responses during a subsequent infection

**DOI:** 10.1101/2022.09.07.507060

**Authors:** Melissa L. Rioux, Anni Ge, Anthony Yourkowski, Magen E. Francis, Mara McNeil, Alaa Selim, Bei Xue, Joseph Darbellay, Alyson A. Kelvin

## Abstract

Our biggest challenge to reducing the burden of seasonal influenza is the constant antigen drift of circulating influenza viruses which then evades the protection of pre-existing immunity. Continual viral infection and influenza vaccination creates a layered immune history in people, however, how host preimmunity interacts with an antigenically divergent virus exposure is poorly understood. Here we investigated the influence of host immune histories on influenza viral mutations. Immune backgrounds were devised in mice similar to what is experienced in people: naive; previously infected (A/FM/1/1947); previously vaccinated (Sanofi quadrivalent vaccine); and previously infected and then vaccinated. Mice were challenged with the heterologous H1N1 strain A/Mexico/4108/2009 to assess protection, viral mutation, and host responses in respect to each immune background by RNAseq. Viral sequences were analyzed for antigenic changes using DiscoTope 2.0 and Immune Epitope Database (IEDB) Analysis Resource NetMHCpan EL 4.1 servers. The mock infected-vaccinated group consistently had the greatest number of viral mutations seen across several viral proteins, HA, NA, NP, and PB1 which was associated with strong antiviral responses and moderate T cell and B cell responses. In contrast, the preimmune-vaccinated mice were not associated with variant emergence and the host profiles were characterized by minimal antiviral immunity but strong T cell, B cell, and NK cell responses. This work suggests that the infection and vaccination history of the host dictates the capacity for viral mutation at infection through immune pressure. These results are important for developing next generation vaccination strategies.

**Importance:** Influenza is a continual public health problem. Due to constant virus circulation and vaccination efforts, people have complex influenza immune histories which may impact the outcome of future infections and vaccinations. How immune histories influence the emergence of new variants and the immune pressure stimulated at exposure is poorly understood. Our study addressed this knowledge gap by utilizing mice that are preimmune to influenza viruses and analyzing host responses as well as viral mutations associated with changes in antigenicity. Importantly, we found previous vaccination induced immune responses with moderate adaptive immunity and strong antiviral immunity which was associated with increased mutations in the influenza virus. Interestingly, animals that were previously infected with a heterologous virus and also vaccinated had robust adaptive responses with little to no antiviral induction which was associated with no emergence of viral variants. These results are important for the design of next generation influenza vaccines.

## Introduction

Despite recurrent vaccination, infections caused by influenza viruses continue to place a heavy burden on public health. Influenza viruses infect an estimated 5-10% of adults and 20-30% of children annually (1). Seasonal influenza viruses specific target the epithelium in the human respiratory tract primarily causing mainly respiratory symptoms of various severity such as coughing, bronchitis or pneumonia which in some cases can lead to death (2). The ultimate outcome of infection depends on a number of host and viral factors (3, 4).

Influenza viruses are negative-sense, single-stranded RNA viruses belonging to the Orthomyxoviridae family of viruses (5). Of the four types of influenza viruses (types A through D), only types A and B circulate seasonally in humans, and only type A has historically caused pandemics (6). Type A influenza viruses are further subtyped by the external and immunologically dominant surface envelop glycoproteins hemagglutinin (HA) and neuraminidase (NA), of which there are 18 HA (H1-H18) and 11 NA (N1-N11) (7). The eight segments of single-stranded, negative-sense RNA encode for at least eleven different viral proteins: HA, NA, matrix 1 (M1), matrix 2 (M2), nucleoprotein (NP), non-structural protein 1 (NSP1), nuclear export protein (NEP), polymerase acidic protein (PA), polymerase basic protein 1 (PB1), and polymerase basic protein 1-F2 (PB1-F2) (8). Genetic variability occurs through two mutational mechanisms: antigenic shift and antigenic drift (5). Antigenic drift is the accumulation and retention of viral mutations during replication which leads to seasonal epidemics, whereas antigenic shift is the swapping of viral genomic segments between different viruses as they infect the same cell at the same time which causes pandemics (5).

Annual vaccination is recognized as the best way to prevent morbidity and mortality from influenza virus infection (9, 10). However, antigenic drift creates the need to reformulate influenza virus vaccines each year leading to annual revaccination of the population (11). Although several vaccine platforms exist, inactivated split virion influenza vaccines are most often used in either a trivalent or quadrivalent formula. Hemagglutinin is the main component of influenza vaccines. It is a homotrimeric protein, and each subunit monomer is made up of two domains referred to as HA1 and HA2 (12). The HA1 domain contains four antigenic sites and the HA1 domain is mostly composed of antiparallel beta-sheets in the globular “head” region. HA2 or the stalk is comprised of three alpha helices, one from each monomer (13). An HA monomer can be further defined by distinct subdomains: the globular head of HA1 contains the N-terminal F’ subdomain (residues 1-41), a vestigial esterase subdomain (residues 42-109 and 263-272), and importantly, the receptor binding domain (RBD) of HA (residues 110-262) (12). The stalk region of HA, which is highly conserved among influenza A viruses, is composed of the F’ (273-330) and F (348-515) subdomains, the fusion peptide (331-347), a transmembrane domain (516-537), and a cytoplasmic domain at the C-terminus of HA2 (538-552)(12). The HA protein is highly dynamic and plastic acquiring the majority of viral mutations of antigenic drift (14). Drifted viruses can antigenically distinct and they can also elicit cross-protective responses if there are epitope similarities in viral proteins of the first and sequential virus (15).

The early stages of influenza virus infection in a naive individual are defined by the antiviral response and innate immunity at the site of infection in respiratory epithelium (16). Viral RNA is detected by pattern recognition receptors (PRRs) such as toll-like receptors TLR3 or 4 or retinoic acid inducible gene I (RIG-I), melanoma differentiation-associated gene 5 (MDA5), and nucleotide-binding oligomerization domain-containing protein 2 (NOD2) which initiates type I and III interferon responses (17-20). Pro-inflammatory cytokines and chemokines such as interleukin (IL)-6, CCL5, CXCL10, IL-8, TNF-a, and CCL2 for leukocyte recruitment and adaptive immune response activation are upregulated early (21-23). Through coordinated efforts with antigen presenting cells (APCs), the adaptive immune responses are initiated to be highly specific to a given epitope on a pathogen (24). In the lymph node, lymphocytes recognize antigens on their T cell receptors (TCRs) or immunoglobulins (Igs) or B Cell Receptors (BCRs) for T cells and B cells, respectively. The APC presents peptides of specific antigen to the T cell through presentation on host major histocompatibility complex (MHC) molecules on the surface of the APC to the T receptor. Engagement of the TCR and MHC initiates cellular activation (25, 26). Together, the MHC molecules (MHC class I (endogenous pathway) and MHC class II (exogenous pathway)) molecules loaded with peptides engage with T cell receptors, CD8+ and CD4+ T cells, respectively (24).

Due to the diversity of influenza viruses and vaccines, humans have unique immunological backgrounds composed of immunity acquired from previous exposures. The continual interactions create of complex web of several host and viral factors that intercept at a viral infection or vaccination event, however, little is understood regarding the contribution of specific pre-existing immunity to future immune responses and how specific immune pressure may influence viral mutation at a new infection. We hypothesize that the unique immune backgrounds will regulate specific immune responses elicited at a new viral exposure event and this will in turn influence viral replication and selection of mutations. Here we address our hypothesis leveraging a C57Bl/6j mouse model of influenza virus infection and vaccination. Immune backgrounds were developed by non-lethal infection with A/FortMonmouth/1/1947 H1N1 virus, vaccination with the 2018-2019 split virion quadrivalent vaccine, and final challenge with an antigenically divergent strain of H1N1 influenza A virus A/Mexico/4108/2009. RNA extracted from lungs collected on day 3 post infection was sequenced for both viral mutations and host expression analysis which identified unique immune responses for each background and specific mutations associated. In particular, naive-vaccinated mice had the most viral mutations with the greatest structural changes across all genes. As these immune background recapitulate the host immune status found within the human population, our study brings essential insight into the mechanisms of the host-virus interaction which is important for the design of next-generation influenza virus vaccines.

## RESULTS

Most people have a complex history of influenza virus infections and vaccinations. Although an individual will have several exposures to influenza viruses by adulthood, influenza viruses continue to evade pre-existing immunity and circulate across the globe (27). Here we designed a study (**Figure 1A**) to investigate both influenza viral mutation and host immune responses following an influenza virus infection in the preimmune host. Immune histories were developed similarly as we had done previously in a ferret study (28). C57Bl/6j mice were infected intranasally with a historical H1N1 strain A/FortMonmouth/1/1947 (FM/47) at 10^3.5^ TCID_50_ on day 0 and left to develop immune memory over two months. Mice were then vaccinated intramuscularly with the Sanofi Fluzone quadrivalent influenza vaccine (QIV) from 2018-2019 on day 60 and boosted on day 74.

**Figure 1.**
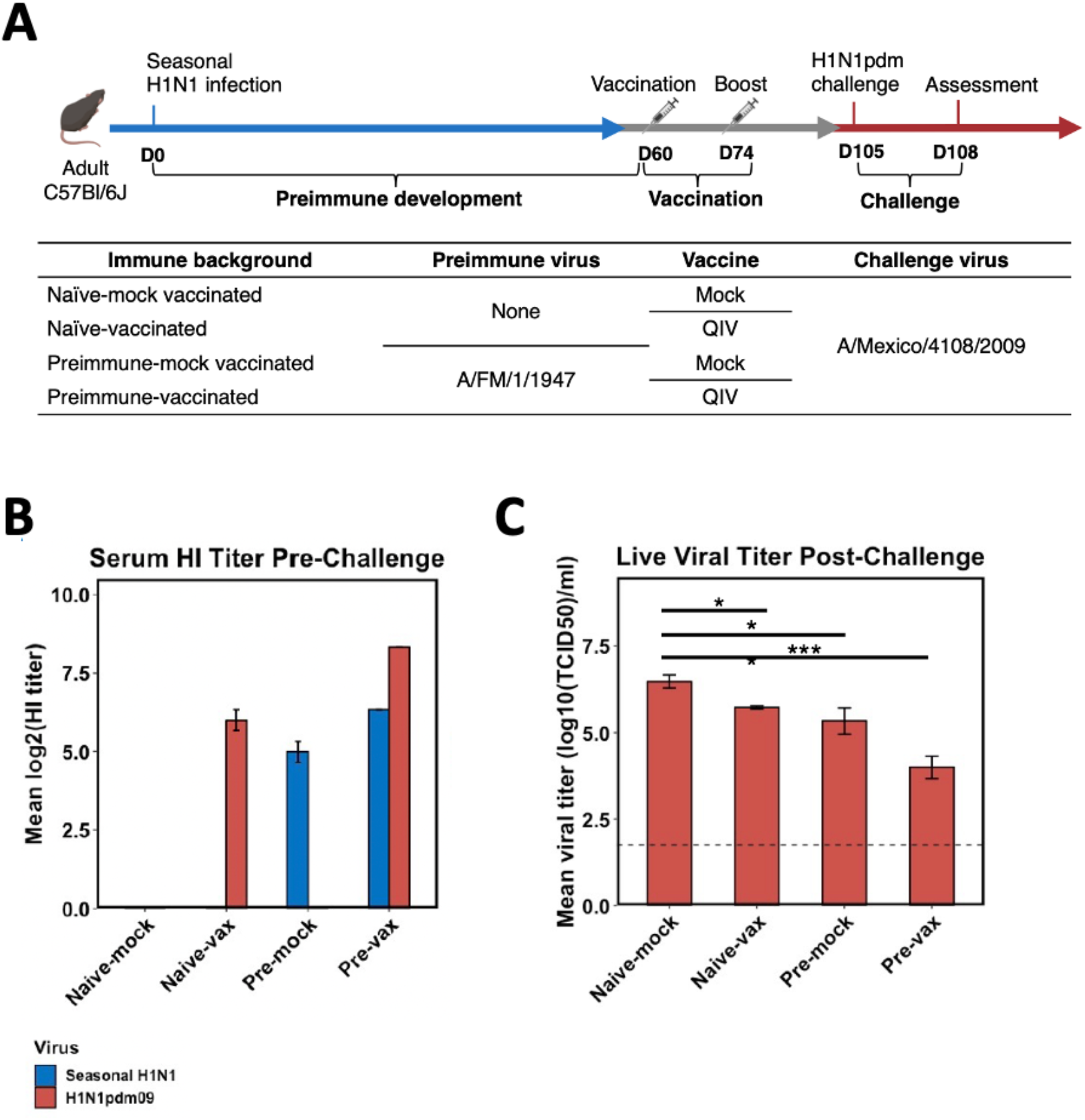
Developing influenza virus immune backgrounds in mice. (A) A schematic of the experimental outline is shown. At day 0, C57Bl/6j mice were intranasally infected with seasonal H1N1 strain A/FortMonmouth/1/1947 (FM/47), followed by intramuscular vaccination and boost with the 2018-2019 quadrivalent vaccine on days 60 and 74. Four immune backgrounds were devised depending on host infection and vaccination status: naïve-mock vaccinated, naïve-vaccinated, preimmune-mock vaccinated, and preimmune-vaccinated. On day 105 of the study, mice of all immune backgrounds were subject to a lethal dose of pandemic H1N1 strain A/Mexico/4108/2009 (Mex/09), followed by assessment of host responses and virus on day 108 (day 3 post challenge (pc)). (B) Three days prior to challenge, antibody titers were quantified against FM/47 and Mex/09 viruses by HAI assay. (C) At three days pc, live viral titer was measured in the lungs. Groups were compared by one-way ANOVA (significance codes: 0 ‘***’ 0.001 ‘**’ 0.01 ‘*’ 0.05 ‘.’ 0.1 ‘’ 1) and Tukey multiple comparisons of means (95% family-wise confidence level).

The vaccine contains HA proteins from the following influenza viruses: H1N1 (A/Michigan/45/2015 X-275) (2009 H1N1 lineage), H3N2 (A/Brisbane/1/2018 X-311), influenza B-Yamagata lineage (B/Phuket/3073/2013), and influenza B-Victoria lineage (B/Colorado/6/2017-like virus). One month after vaccination, mice were challenged intranasally with a lethal dose of A/Mexico/4108/2009 (Mex/09) at 10^6^ TCID50 and assessed at three days post-challenge (29). Importantly, the challenge virus was antigenically similar to the H1 vaccine component of the vaccine. The imprinting virus FM/47 is antigenically distinct from the challenge Mex/09 with 78% identity in the HA protein. Four experimental groups were designed as outlined in **Figure 1A** to control the study at challenge: naïve-mock vaccinated (no pre-existing immunity), naïve-vaccinated (only vaccinated), preimmune-mock vaccinated (only previously infected with FM/47), and preimmune-vaccinated (both previously infection and vaccinated). The immune status of the mice was confirmed by hemagglutination inhibition (HI) assay against preimmune (FM/47) and vaccine (Mex/09) influenza strains (**Figure 1B**) prior to challenge. Confirming vaccination, naïve-vaccinated and preimmune-vaccinated mice had average log2(HI titers) of 6 and 8 HAI units, respectively, toward Mex/09. Both preimmune groups had high HI titer against the preimmune FM/47 strain (5 and 7 HAI units in preimmune-mock and preimmune-vaccinated groups, respectively). Following Mex/09 challenge, lung tissue was collected from mice on day 3 pc (post challenge) for assessment of viral load as well as downstream sequencing analysis for viral mutations and host responses (workflow illustrated in **Figure 2**). Importantly, live virus was detected in the lungs of mice from all immune backgrounds, however, the levels of the virus was associated with the layering of previous immunity as the naive-mock vaccinated mice had the highest lung viral titer at ∼6 log_10_TCID_50_ml^-1^ and the preimmune-vaccinated mice had the lowest at ∼4 log_10_TCID_50_^ml-1^ (**Figure 1C**). The viral load in all groups was statistically different compared to the naive-mock group. These data suggested that despite a layered immune background that a subset of the initial viral population is able to evade sterilizing immunity in mice with pre-existing immunity.

**Figure 2.**
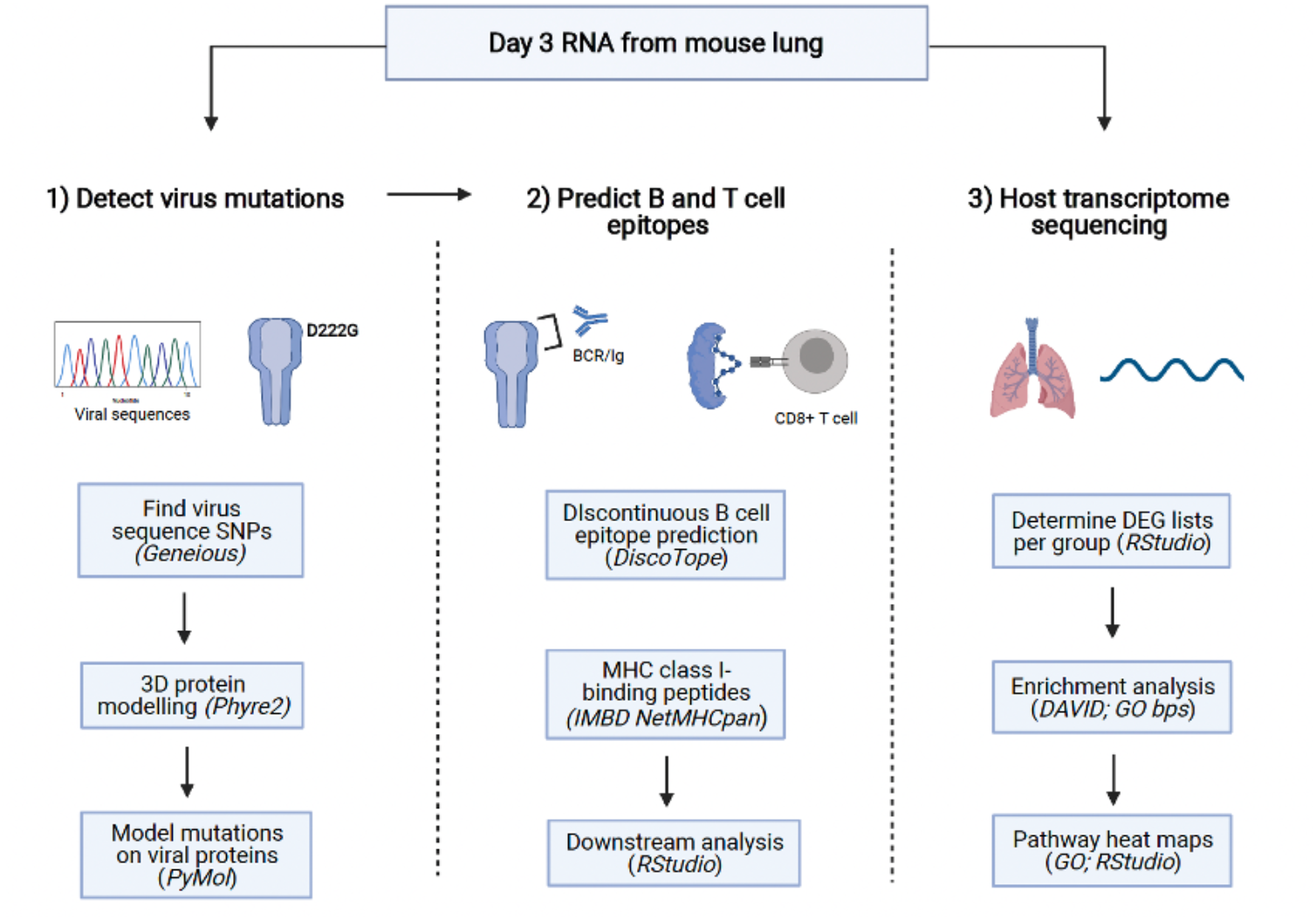
Schematic of the bioinformatic analysis pipeline used to determine influenza virus mutations, predicted B and T cell epitopes, and host transcriptome analysis after challenge. 1.) At three days pc with A/Mexico/4108/2009 (Mex/09), viral RNA from mouse lungs was sequenced using Illumina MiSeq. Viral sequences were aligned to reference Mex/09 using Geneious, and SNPs and variants above 1% frequency were detected for sequences extracted from mice of each immune background and aligned using MEGAX. To predict the possible structural and antigenic impact of immune background specific amino substitutions, representative 3D models were generated using the Phyre2 platform and images were generated in PyMol. 2.) The 3D model was used to predict discontinuous B cell epitopes along the viral proteins of each group using DiscoTope 2.0 (DTU Health Tech) and variant amino acid sequences were uploaded to the Immune Epitope Database NetMHCpan EL 4.1 tool to predict T cell epitopes. 3.) The host transcriptional response was investigated by sequencing of mouse lung RNA. Lists of significantly differentially expressed genes (DEGs) for each immune background were uploaded to the Database for Annotation, Visualization and Integrated Discovery (DAVID) version 6.8 functional annotation tool to determine enriched immune pathways. Heatmaps were generated in for antiviral, T cell, and B cell-mediated immune response pathways.

### Combined preimmunity and vaccination decreases the frequency and number of influenza virus single nucleotide polymorphisms post-challenge

Since viral replication was observed in the lungs of preimmune-vaccinated mice as well as in the mice of other immune backgrounds, we next characterized the viral population that had evaded pre-existing immunity. To investigate the viral quasispecies and variants that present in the mice during infection, Illumina MiSeq whole-viral genome sequencing was performed on viral RNA isolated from lung tissue at three days post-challenge as well as the Mex/09 virus stock for comparison (**Figure 2**). The viral sequences are available on the Sequence Read Archive (SRA) and reads were aligned to Mex/09 reference sequences for quantification (**Table 1**) using Geneious software (30). The sequenced stock Mex/09 yielded the highest number of reads and coverage across most segments compared to virus extracted from mouse lungs, as expected.

**Table 1.** Mean number of reads and coverage per influenza virus gene following alignment of viral sequences to A/Mexico/4108/2009 reference segments in Geneious.

| Gene | Length (nt) | Stock Mex/09 |  | Naïve-mock vaccinated |  | Naïve-vaccinated |  | Preimmune-mock vaccinated |  | Preimmune-vaccinated |  |
| --- | --- | --- | --- | --- | --- | --- | --- | --- | --- | --- | --- |
|  |  | Mean reads | Mean coverage | Mean reads | Mean coverage | Mean reads | Mean coverage | Mean reads | Mean coverage | Mean reads | Mean coverage |
| PB2 | 2280 | 152,153 | 36,192 | 116,243 | 18,782 | 144,600 | 22,568 | 156,061 | 19,354 | 201,613 | 26,076 |
| PB1 | 2274 | 111,003 | 18,718 | 78,144 | 10,244 | 122,704 | 17,736 | 94,685 | 9378 | 92,614 | 8949 |
| PA | 2151 | 75,936 | 13,124 | 52,570 | 5553 | 19,548 | 1564 | 46,844 | 3716 | 54,251 | 4773 |
| HA | 1701 | 42,205 | 5111 | 6787 | 799 | 11,001 | 1091 | 9939 | 899 | 5776 | 529 |
| NP | 1497 | 6591 | 771 | 1831 | 206 | 632 | 74 | 2249 | 185 | 1375 | 140 |
| NA | 1410 | 20,003 | 2382 | 1430 | 169 | 1904 | 119 | 1027 | 130 | 2935 | 410 |
| M1, M2 | 982 | 43,187 | 7426 | 11,507 | 2095 | 5142 | 942 | 7939 | 1315 | 13,928 | 2335 |
| NS1 | 844 | 60,291 | 11,449 | 16,024 | 3330 | 14,358 | 2999 | 17,260 | 3364 | 47,157 | 6245 |

We next analyzed the mutations within the sequencing reads to determine if novel viral mutations had been acquired in influenza virus genes extracted from mouse lungs, or if variants already present in the stock virus had changed in frequency. The insertions, deletions, and single nucleotide polymorphisms (SNPs) above one percent frequency (minimum of five reads) present on the gene sequences were determined using the SNPs/variants tool in Geneious (as outlined in **Figure 2**). Manhattan plots and frequency histograms were generated to visualize the distribution of viral SNPs and their frequencies across groups and over viral genes (**Figure 3**). Highly significant (p << 0.05) SNPs were detected across all ten influenza virus genes and across all immune backgrounds, except for M1/M2 segment of the stock virus, and M2 gene in the naïve-mock vaccinated group. For HA, naïve-mock vaccinated mice had markedly less SNPs than the stock virus used to inoculate animals, indicating selection of variants adaptive to the mouse model. More mutations were detected in naïve-vaccinated mice in HA compared to all other groups, with an increased number of SNPs that were highly significant (p << 0.05). Although preimmune groups had a lower number of detectable SNPs across HA, a greater proportion of these SNPs were highly significant compared to the HA sequences from naïve mice. The trend of increased detectable SNPs in naive-vaccinated mice was generally consistent for the other influenza virus genes. Interestingly, the groups with only one exposure, the naive-vaccinated and preimmune-mock vaccinated groups, had the greatest number of mutations in the Nucleocapsid protein (NP) and neuraminidase (NA) while the least detected was in preimmune-vaccinated animals (**Figure 3A**).

**Figure 3.**
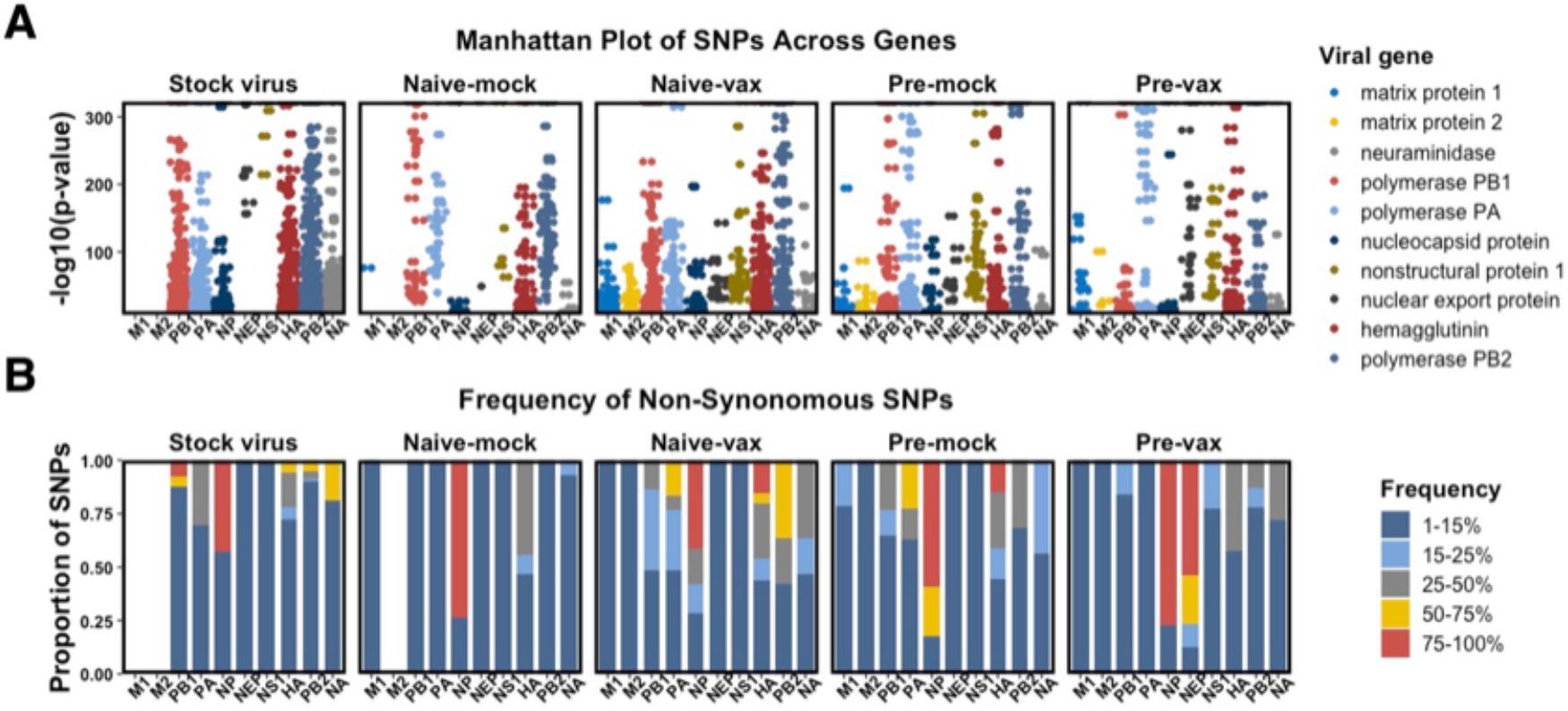
Influenza virus gene sequences extracted from mice of different immune backgrounds pc have significant SNPs on viral genes analyzed. Viral sequences were aligned to all reference Mex/09 genes (accession numbers as follows: PB2 (GQ379815), PB1 (GQ149652), PA (GQ149653), HA (GQ223112), NP (GQ149655), NA (GQ149650), M2 (GQ149657), and NEP (GQ149658) using Geneious, and SNPs and variants above 1% frequency were detected for and plotted per immune background. Stock Mex/09 used to inoculate animals was sequenced and compared. (A) The SNPs found in all ten influenza virus genes were visualized using a Manhattan plot to determine the number and significance (-log10(p-value) of each mutation. (B) After translating the genetic sequence to protein sequence, the proportion of non-synonymous SNPs detected for each protein were categorized by their frequency, with frequency bins as follows: 1-15%, 15-25%, 25-50%, 50-75%, and 75-100%.

We next examined the frequency distribution of SNPs across viral genes leading to amino acid changes. Substitution, truncation, or start codon loss across viral genes were then represented on histograms after the frequency was binned (1-15%, 15-25%, 25-50%, 50-75%, and 75-100%) (**Figure 3B**). The M1/M1 segment was interesting as there were no SNPs of any frequency detected in the stock virus or in naïve-mock vaccinated animals, however, SNPs appeared on M1/M2 in naïve-vaccinated, preimmune-mock vaccinated, and preimmune-vaccinated groups after challenge (**Figure 3B**). For HA sequences, at least one quarter of the total SNPs fell into the higher-frequency ranges (> 15%) in all experimental groups with over half of all HA SNPs in the sequences from the naive-mock and naive-vaccinated animals occurring above 15% frequency. The HA analysis indicated that the presence of previous infection combined with vaccination, or conversely, no previous immunity against the challenge strain, was associated with a decrease in high-frequency variants. Overall, this preliminary analysis suggests a relationship between the infection and vaccination history of the host and potential trends for viral mutation in terms of significance and SNP frequency.

### The hemagglutinin protein shows immune background-specific mutations that may impact protein structure

We next focused our analysis on the HA protein due to its role in viral entry, immunogenic nature, and propensity for mutation (31). SNPs that led to non-synonymous changes were plotted along the amino acid position in the HA amino acid sequence (**Figure 4A**). The stock Mex/09 virus used to inoculate animals had the greatest number of HA variants which decreased in all immune backgrounds after replication. Novel mutations not originally detected in stock virus also emerged in naïve-mock vaccinated animals, suggesting the emergence due to errors in replication.

**Figure 4.**
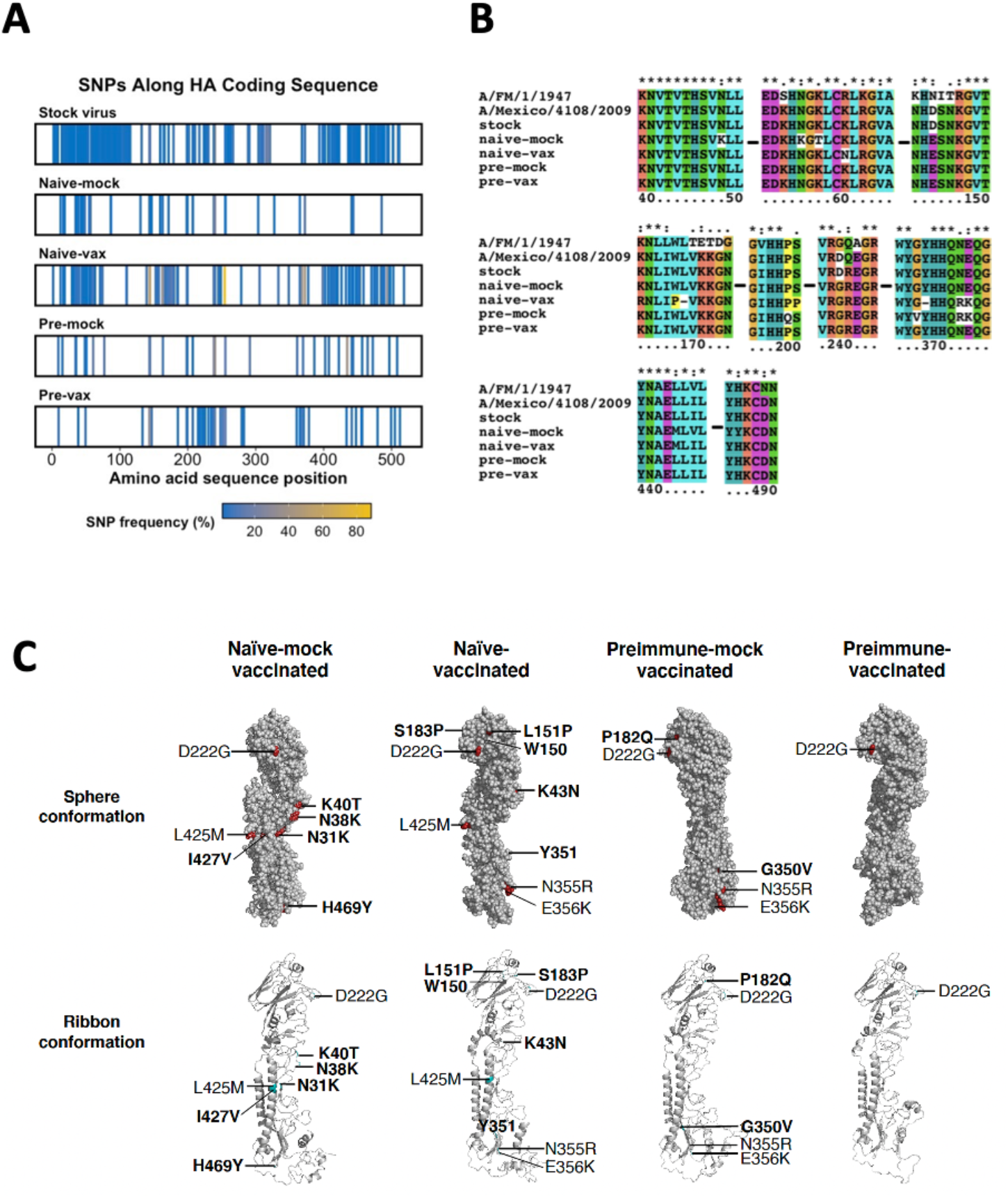
Naïve-vaccinated animals have the highest frequency of SNPs across influenza virus HA pc. (A) The non-synonymous SNPs found in HA gene sequences after introduction to naïve-mock vaccinated, naïve-vaccinated, preimmune-mock vaccinated, and preimmune-vaccinated mice were plotted on the amino acid sequence number of HA (1 band = 1 SNP) generating a bar-code effect for each SNP profile. The color of the band represents the relative frequency of each mutation. (B) HA gene sequences from mice of each immune background were aligned using MEGAX. The amino acid sequences containing immune background-specific substitutions were shown using ClustalX in the alignment. (C) To predict the possible structural and downstream antigenic impact of each immune background-specific amino substitution, a representative 3D model of each HA protein was generated using the Phyre2 platform and protein images were generated in PyMol to show both sphere (top) and ribbon (bottom) conformations. Group-specific SNPs are highlighted in bold and non-bolded text represents SNPs not unique to a group.

Of the four immune backgrounds, the greatest number of SNPs identified on HA post-challenge were detected in naïve-vaccinated animals, with the majority of the mutations being in the 1-20% frequency range (**Figure 3B**; **Figure 4A**). In this group the highest density of HA mutations were found in the N-terminal F’ subdomain (amino acids 1-41, H1 numbering), the RBD (110-262), and the F subdomain in the stalk region (348-515) (**Figure 4A**) (12). Preimmune-mock vaccinated animals had a relatively even distribution of mutations across the entire HA protein, while mutations in preimmune-vaccinated mice were most dense in the RBD (**Figure 4A**). To compare the shared amino acid changes among in each experimental group, the HA sequences were aligned using MEGA (32) (**Figure 4B**). Interestingly, the number of shared mutations between animals of a group was not associated with the overall number of SNPs that persisted in that group (**Figure 4B**; summarized in **Table 2**).

**Table 2.**
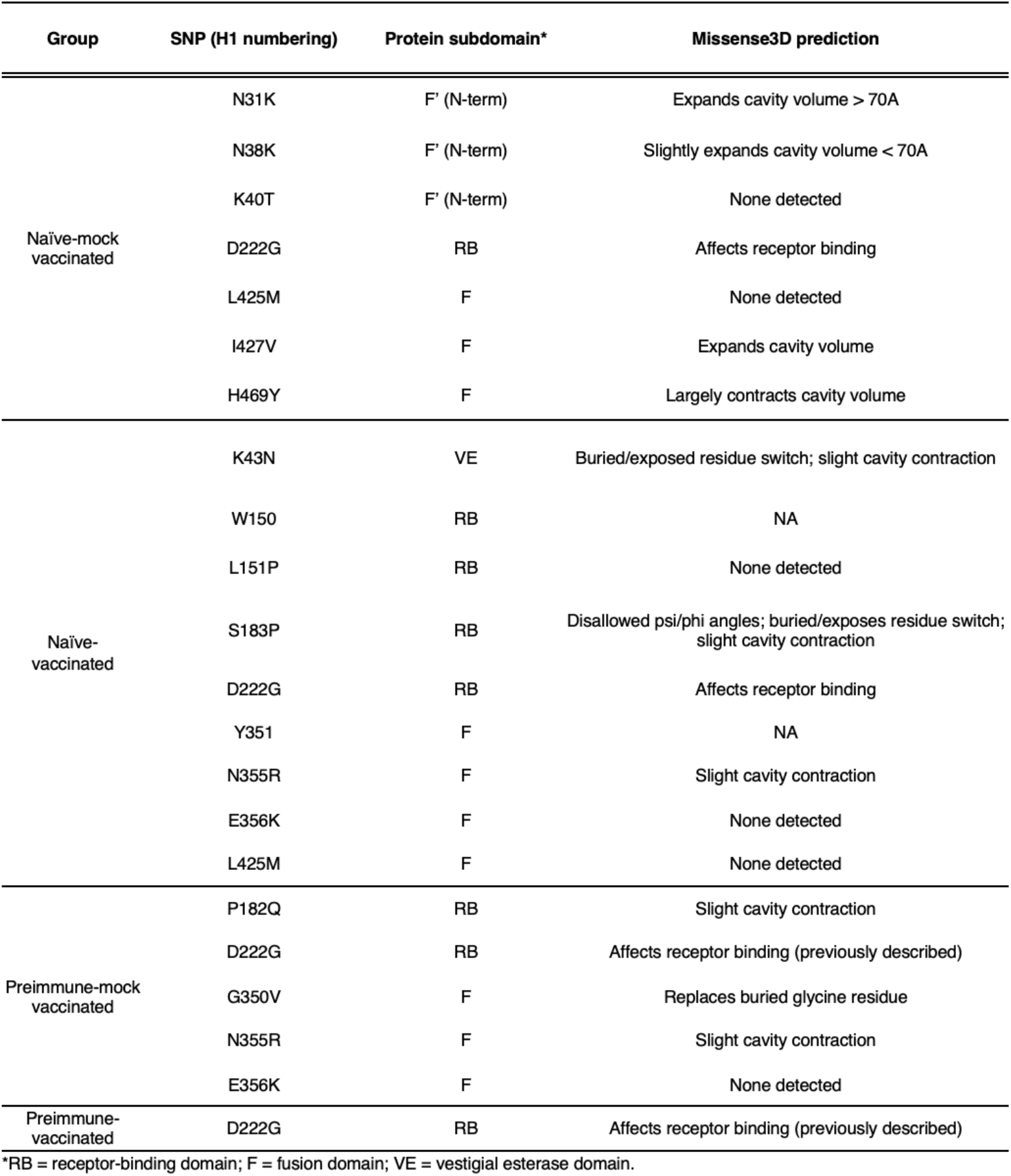
Summary of immune background-specific single nucleotide polymorphisms on influenza virus hemagglutinin and potential structural damage predicted by Missense3D.

To determine how each of these mutations may impact protein structure and antigenicity, protein folding predictions were generated using the Phyre2 platform and PyMOL (The PyMOL Molecular Graphics System, Version 2.3.5, Schrödinger, LLC) was used to generate a representative 3D model (**Figure 4C**) (33). The mutation D222G in the RBD of HA was detected in mice of all immune backgrounds and was the only common mutation identified from the preimmune-vaccinated mice (**Figure 4c**). For each of the remaining immune backgrounds, group-specific mutations were identified in both the head and stalk regions of HA (**Table 2** and bolded in **Figure 4C**). Naïve-mock vaccinated mice had several group-specific mutations in the stalk region of the protein, including N31K, N38K, K40T, I427V, and H469Y, as well as L425M shared with naïve-vaccinated mice (**Table 2**). Group-specific mutations were also detected in naïve-vaccinated mice, such as K43N (stalk) as well as three mutations (W150, L151P, and S183P) in the RBD (**Figure 4C**; **Table 2**). Finally, preimmune-mock vaccinated mice shared only P182Q in the RBD and G350V in the protein stalk (**Figure 4C**; **Table 2**). The potential effect on protein structure of each amino acid substitution was analyzed using Missense3D which identified possible predicted structural damage (summarized in **Table 2**) (34). Some notable mutations included the HA stalk H469Y identified in naïve-mock vaccinated mice suggested a contraction (>70Å) of the cavity as well as two substitutions in the F’ subdomain (N31K, N38K) and one in the fusion subdomain (I427V) each predicted to expand the cavity volume (**Table 2**). Of the nine substitutions associated with naïve-vaccinated animals post-challenge, three were predicted to cause structural impacts: K43N in the vestigial esterase domain and; S183P in the RBD predicted to cause a buried residues to become exposed and N355R suggesting a cavity contraction < 70Å (**Table 2**) (12). Finally, three of the five substitutions found in preimmune-mock vaccinated animals had reported structural damage: one in the receptor-binding domain (P182Q) (cavity contraction < 70Å) and two substitutions G350V and N355R (**Table 2**). Similar to previous assessments of impacting mutations, naïve-vaccinated and preimmune-mock vaccinated animals had the greatest number with the least in the mice with layered or naive immune backgrounds.

### Immune background-specific mutations in hemagglutinin alter predicted antigenicity

Amino acid mutations can lead to changes in antigenicity with respect to B cell epitopes and the specific peptides presented to T cells. Peptide presentation of antigen by antigen presenting cells or infected cells is essential for efficient and accurate T and B cell activation and maturation. T cell epitopes are linear peptides, consisting of approximately 8-11 amino acids for MHC class I, while B cell epitopes typically require a 3D confirmation of continuous or discontinuous amino acids for recognition (35). Since both arms of the immune system are important for clearing a pathogen and establishing immune memory, the predicted B cell and T cell epitopes for the HA molecules sequenced from each immune background were assessed.

Starting with the group-specific HA folding models generated by Phyre2, the discontinuous B cell epitopes per immune background were mapped using DiscoTope 2.0 server (threshold of −3.7 (0.47 sensitivity, 0.75 sensitivity)) (35) (**Figure 5A** and **B**). Peaks appearing above and valleys below the prediction threshold indicate regions that are likely and unlikely to be B cell epitopes, respectively. Marked differences in the predicted B cell epitopes, both in the “size” of the epitope (i.e., the number of amino acids included in an epitope), and in the DiscoTope score (i.e., propensity to be a B cell epitope) were identified when sequences associated with each immune background were compared to the reference sequence. Differences in predicted B cell epitopes were evident in both the head (amino acids 42-272) and stalk (331-552) regions of the HA protein (**Figure 5C**). The epitope appearing at residues 137-147 in the RBD of HA scored more highly in the naïve-vaccinated group compared to the Mex/09 reference or the other immune backgrounds (**Figure 5B**). Residues 197-215 and 227-241 in the RBD had increased DiscoTope scores in naïve-vaccinated mice compared to the other sequences. Another small epitope occurring at residue 261 was also apparent in HA from naïve-vaccinated animals only (**Figure 5B**). Other predicted epitope changes occurred outside of the RBD of HA such as at residues 405-416 in the fusion subdomain in sequences from naïve-vaccinated, preimmune-mock vaccinated, and preimmune-vaccinated groups. Another predicted epitope region in the fusion subdomain (residues 496-505) was expanded in the preimmune-vaccinated group (**Figure 5B**). Taken together, our analysis of predicted B cell epitopes indicated that immune background-specific mutations occurring on the HA protein after challenge may lead to antigenic differences in replicating viruses which were again most prominent in the HA of naïve-vaccinated animals.

**Figure 5.**
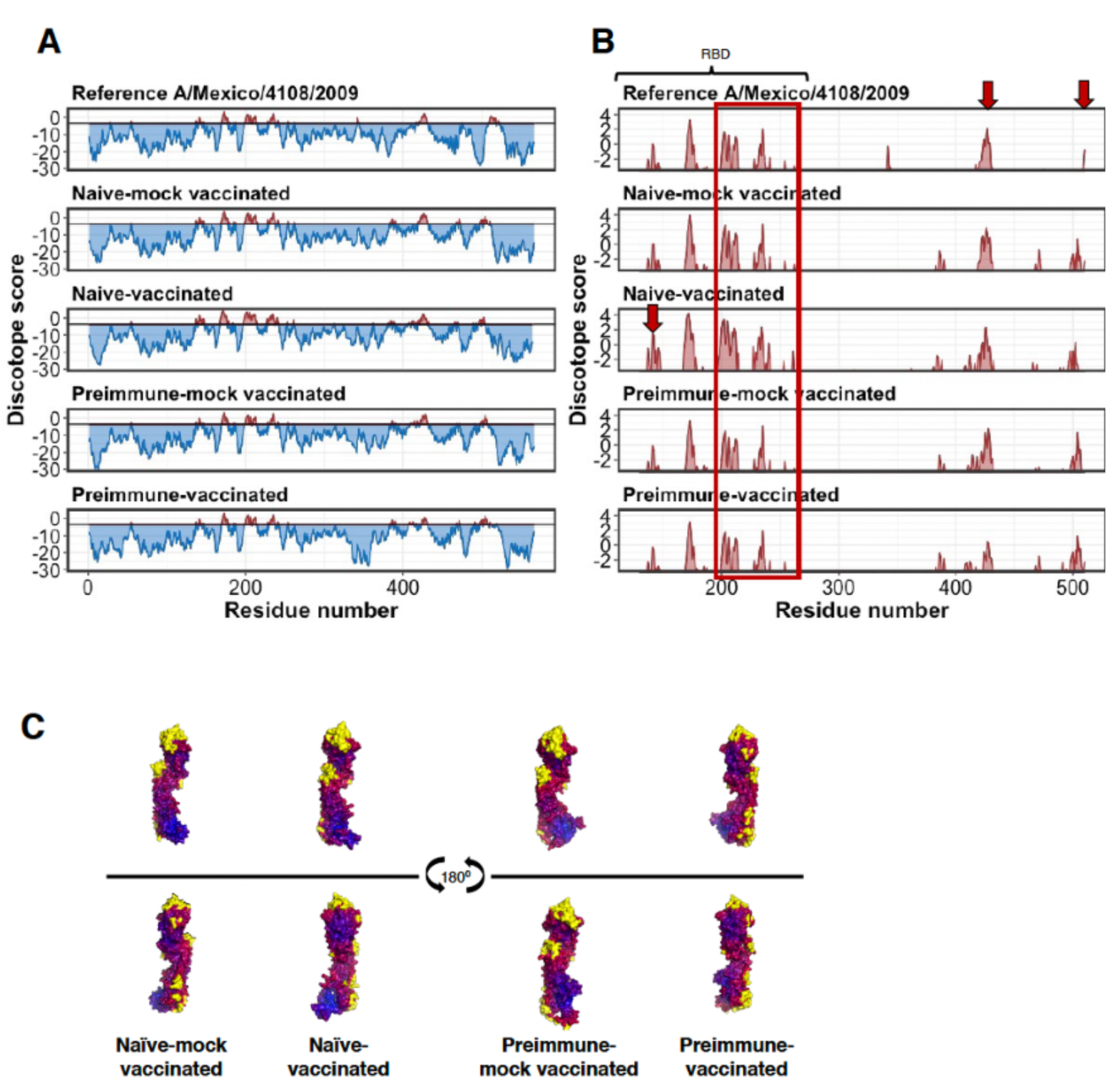
Influenza virus HA proteins show changes in predicted B cell epitopes after viral replication in mice with varying immune backgrounds. The folding models of HA generated with the Phyre2 platform were used to identify HA surface epitopes using DiscoTope 2.0. (A) DiscoTope scores falling below (blue) and above (red) the B cell epitope prediction threshold of −3.7 (0.47 sensitivity, 0.75 specificity) were mapped against amino acid sequence number for reference Mex/09 (NCBI accession no. GQ223112) and the sequences acquired in mice of varying immune backgrounds at challenge. (B) For greater resolution, only the positive B cell epitope prediction results are shown. The arrow highlights notable differences in DiscoTope scores compared to reference Mex/09, which fall outside of the RBD of HA. The red square denotes areas of notable variation among sequences. (C) Representative folding models of the HA protein showing predicted B cell epitopes after challenge in mice of different immune backgrounds were made in PyMol and DiscoTope scores are shown as heatmaps along the folded protein, with residues colored according to their predicted score: yellow indicates positively predicted B cell epitopes (scores > −3.7 threshold), red indicates high-scoring amino acids, and blue indicates low scoring regions (i.e., unlikely B cell epitopes).

We next evaluated the impact of mutations on MHC class I presentation for CD8+ T cell activation. Using IEDB NetMHCpan EL 4.1, analysis was restricted to H2-Db and H2-Kb murine alleles and the highest scoring peptides were predicted from our sequences (summarized in **Table 3**). There were several high-affinity, immune-background specific peptides identified across the HA protein that were not originally identified in the stock Mex/09 virus. For example, the peptide LVLLENERTL from amino acids 443-442 (stalk region) was specific to viral sequences from naïve-mock vaccinated animals, while the peptide NAEMLILLENERTL (437-450) spanning the same region of HA scored more highly and was specific to naïve-vaccinated animals (**Table 3**). Additionally, the viral sequences emerging from naïve-vaccinated animals contained several novel peptides in the RBD of HA (specifically 157-168) that were not identified in the other immune backgrounds (**Table 3**). Unlike our previous analysis, the greatest number of high-affinity, group-specific peptides were found in viral sequences isolated from naïve-mock vaccinated animals, with the least found from both preimmune groups.

**Table 3.**
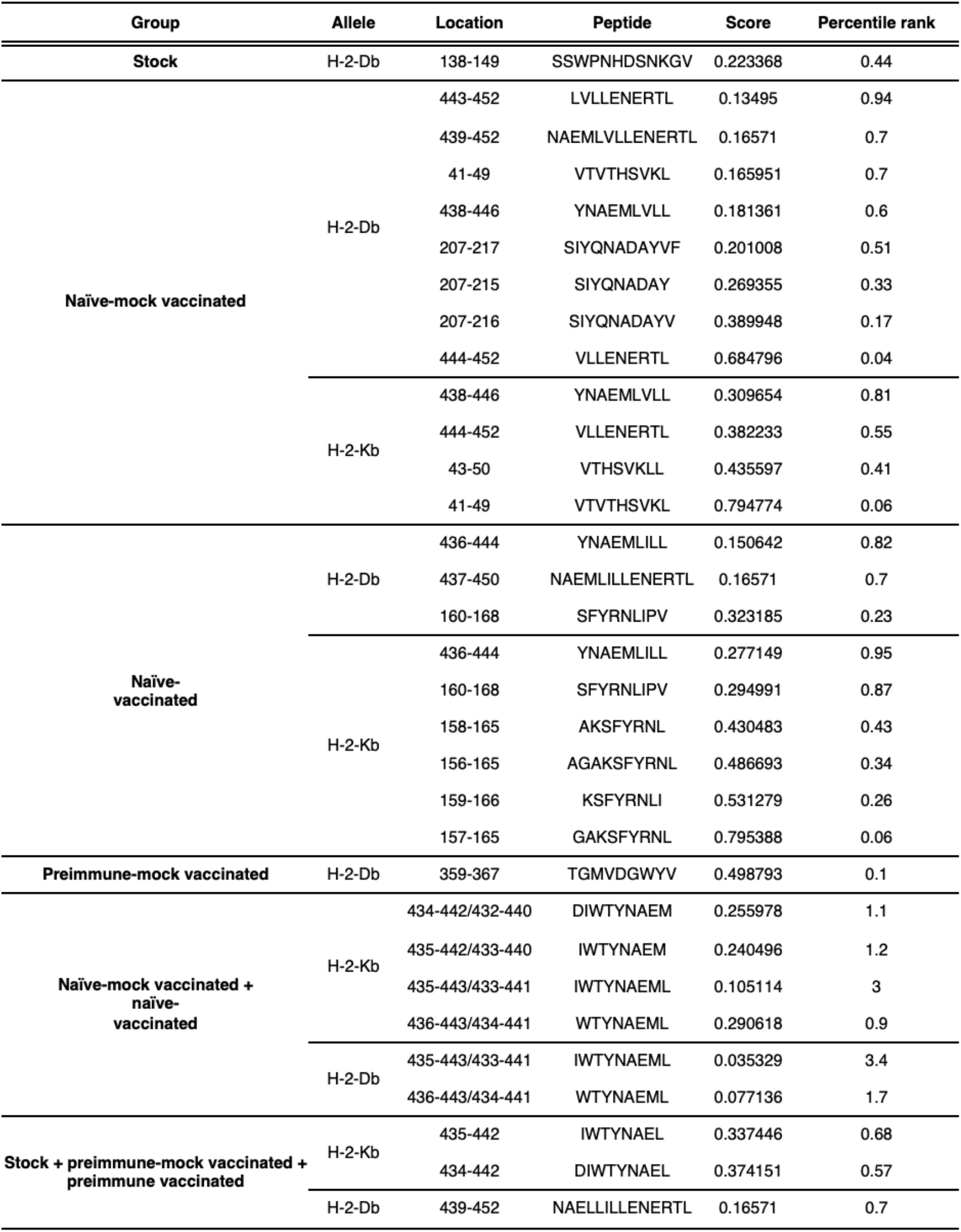
High-affinity MHC class l-binding peptides derived from immune background specific hemagglutinin protein sequences using NetMHCpan EL 4.1.

### Immune background-specific mutations also occur on influenza viral proteins nucleoprotein (NP), neuraminidase (NA), and polymerase basic 1 (PB1)

Although HA is the immunodominant viral protein, other proteins on the surface as well as internal to the virion can contribute to virus antigenicity and have been identified in homologous and heterologous immune responses (36-42). Because these proteins are potential antigenic targets during an influenza virus infection, we also analyzed mutations acquired focusing on the nucleocapsid protein (NP), which is internal and highly conserved; neuraminidase (NA), an external major antigen; and polymerase basic 1 (PB1), a highly conserved polymerase subunit (43).

The acquired viral sequences were mapped to the NP, NA, and PB1 genes to identify all SNPs occurring above 1% frequency. All non-synonymous SNPs found in NP, NA, and PB1 were plotted along the amino acid sequence number of each protein to determine patterns across groups (**Figure S1A**). With respect to the number of SNPs detected across immune backgrounds, NP showed a similar trend to HA, with the greatest number of SNPs found in naïve-vaccinated animals, and the least in preimmune-vaccinated animals (**Figure S1A**). In contrast, NA had the most detected mutations in naïve-mock vaccinated animals, and PB1 mutations were most abundant in naïve-vaccinated and preimmune-mock vaccinated groups (**Figure S1A**). Structural impact analysis identified mutations of high consequence in the NP gene to be associated with naïve-vaccinated animals (**Table S1** and **Figure S1B**). Several specific mutations across NA, including E47G and Q51E (N1 numbering), were located in the stalk domain of sequences from naïve-mock vaccinated and challenged mice (**Table S1**). Additionally, a V291E mutation was identified in the head domain of NA extracted from naïve-vaccinated animals, and I38T in the stalk domain of NA from preimmune-mock vaccinated animals (**Figure S1B**; **Table S1**). Apart from preimmune-vaccinated mice, all groups shared NA sequences with a C to S substitution at amino acid 292 in the head domain which was predicted to induce disulfide bond breakage. Interestingly, only PB1 sequences extracted from preimmune-vaccinated mice had mutations evenly distributed across the entire sequence. The only mutations shared per group for PB1 were detected in the naïve-vaccinated animals (**Figure S1B**). These mutations occurred in several functional domains of the protein, including L95P in the cRNA promoter binding site; M246I in the putative nucleotide binding site; and D617A in the vRNA promoter binding site which were associated with structural alterations that may affect function (**Figure S1B**; **Table S1**) (44). When B cell epitopes were plotted at low and high resolution (**Figure S2A** and **S2B**, respectively) with structural heat maps (**Figure S2C)** the results indicated clear differences in the height and width of peaks corresponding to predicted B cell epitopes in naïve-vaccinated and preimmune-mock vaccinated hosts, specifically at residues 0-25 and 450-500 for NP. Minimal differences in predicted B cell epitopes were observed for NA. No differences were predicted in B cell epitopes for PB1. For CD8+ T cell receptor epitopes presented on H2-Db and H2-Kb alleles we found several group-specific peptides across NP, NA, and PB1, suggesting that variant amino acid sequences may be presented on MHC I molecules and differentially activate CD8+ T cells, giving the variant proteins a unique antigenic presentation.

### Transcriptomic analysis of host immunity in the lung identified specific host responses per immune background that may drive viral mutation at challenge

To complement the viral mutational analysis, we next characterized the host immune response in the lungs during challenge to identify the immune pressure exerted by each pre-existing immune background. We hypothesized that analysis of the immune responses may give insight into the origins of the mutations identified in the viral proteins of host with specific immune backgrounds. To this end, RNA sequencing using the Illlumina NovaSeq 6000 Sequencing System at Novogene (Sacramento, CA, USA) was performed on host lung RNA extracted three days post-challenge. Lung gene expression fold-change was quantified in terms of fragments per kilobases mapped (FPKM) to Mus musculus mm10 exons. The significant (p-value < 0.05) differentially expressed genes (DEGs) between the four immune backgrounds were compared to assess broad similarities and/or differences in the immune response profiles as per numbers of shared genes expressed (**Figure 6A**). At three days post challenge, 625 genes were significantly up- or down-regulated, across immune background, as shown in the Venn diagram (**Figure 6A**). As well, mice of each immune background had significant DEGs that were not shared with the other groups, with 227 DEGs in naïve-mock vaccinated, 1822 in naïve-vaccinated, 305 in preimmune-mock vaccinated, and 872 in preimmune-vaccinated mice (**Figure 6A**). Overall, differential gene expression at challenge was least observed in naïve-mock vaccinated animals, with only 1621 total DEGs detected, compared to 3000-4600 DEGs in the other groups (**Figure 6A**). However, the majority of these DEGs (78%) were also differentially expressed in naïve-vaccinated animals, indicating a high degree of similarity in the immune response profiles, despite a large difference in the number of DEGs overall. In contrast, only 48% of the DEGs found in naïve-mock vaccinated animals were also found in preimmune-vaccinated mice. Naïve-vaccinated mice had the greatest differential gene expression at challenge, with 4680 genes differentially regulated relative to the baseline. When comparing the DEGs present in naïve-vaccinated animals to those found in thev other immune backgrounds, naïve-vaccinated mice were most similar to preimmune-mock vaccinated animals, with 2413 (52%) of the DEGs shared between both groups.

**Figure 6.**
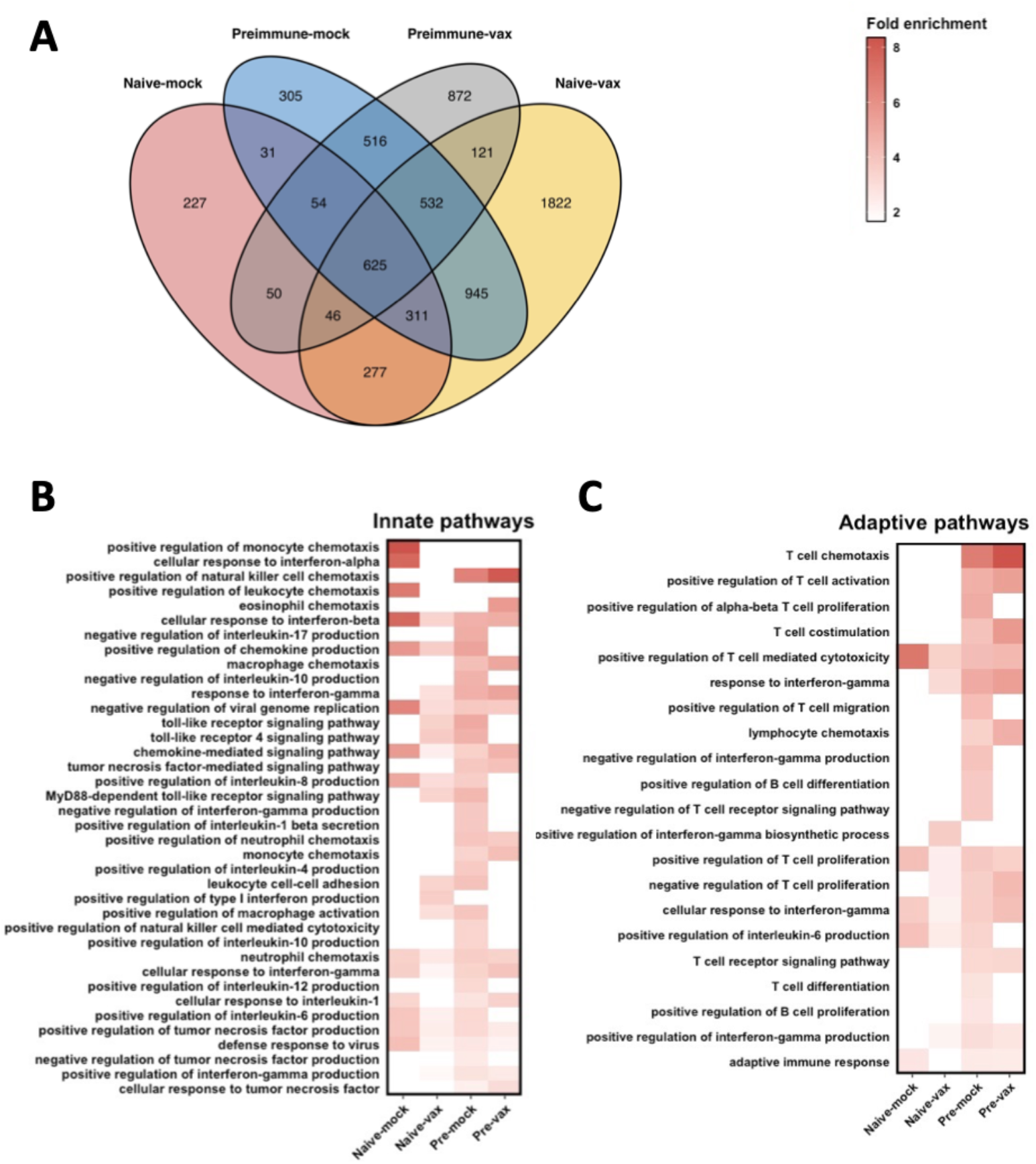
Lungs of mice of different immune backgrounds have different host response gene enrichment profiles at three dc. At three days pc, total RNA was extracted from mouse lungs and the host transcripts were sequenced using the Illumina platform and quantified as fragments per kilobases mapped (fpkm), and the log2 (fold change) was calculated relative to non-infected mice. (A) Similarities and differences in gene regulation between groups are shown in a Venn diagram generated in R using the VennDiagram package version 1.6.20. The lists of differentially regulated genes for each immune background were uploaded to The Database for Annotation, Visualization and Integrated Discovery (DAVID) version 6.8 functional annotation tool for enrichment. GO biological processes were loosely separated into (B) “innate” or (C) “adaptive” immune pathways.

To characterize the dominant immune response pathways of each group at challenge, enrichment analysis by Gene Ontology (GO) in The Database for Annotation, Visualization, and Integrated Discovery (DAVID, https://david.ncifcrf.gov/home.jsp) was used to identify innate (**Figure 6B**) or adaptive (**Figure 6C**) immune signatures. With respect to innate signaling, naïve-mock vaccinated mice had a strong proinflammatory and antiviral response at challenge, highly enriched by monocyte chemotaxis, chemokine (e.g., IL-8) signaling, proinflammatory signaling (e.g., IL-1β and IL-6), and type I and type II interferon response genes (**Figure 6B**). Naïve-vaccinated mice also had enrichment of antiviral (type 1 interferons) and inflammatory response genes and pathways (IL-8, IL-6, and TNF), marked by chemotaxis and cell adhesion (**Figure 6B**). However, naïve-vaccinated and preimmune-vaccinated mice were also enriched in toll-like receptor (TLR) signaling and MyD88-dependent TLR signaling pathways, which were not observed in the naïve-mock vaccinated. Preimmune mice had several commonalities as well as key differences in innate immune involvement at challenge compared to naïve-mock and naïve-vaccinated groups. As found in the naïve animals, preimmune mice also had enrichment of IFN-β and -γ production, leukocyte chemotaxis and cell-cell adhesion, macrophage activation, and TNF production (**Figure 6B**). Differences among the groups included additional enriched pathways such as positive regulation of natural killer (NK) cell activation, IL-1β, and IL-17 were noted, with concomitant negative regulation of IL-10 in preimmune-mock vaccinated mice (**Figure 6B**). Also, preimmune-vaccinated mice had distinct profiles from the other groups marked by monocyte, macrophage, neutrophil, and eosinophil chemotaxis gene enrichment (**Figure 6B**). Enrichment of genes for adaptive immune pathways at challenge was minimal in naïve-mock vaccinated and naive-vaccinated animals (**Figure 6C**). This involvement was restricted to positive regulation of T cell-mediated cytotoxicity and T cell proliferation transcripts in these groups. In contrast, preimmune-mock vaccinated mice had widespread involvement of adaptive immunity (**Figure 6C**) with notable enrichment of lymphocyte chemotaxis, B and T cell differentiation and proliferation, and T cell-mediated cytotoxicity. Additionally, genes for cytokines regulating adaptive immune responses such as IL-12 and IL-4 were also enriched in this group (**Figure 6B** and **6C**). Preimmune-vaccinated mice had adaptive gene regulation that was less diverse than the other experimental groups, but more highly enriched (**Figure 6C**) which included strong enrichment of lymphocyte chemotaxis, positive regulation of T cell activation (TCR signaling and co-stimulation) and cytotoxicity, and regulation of T cell proliferation (**Figure 6C**).

To further characterize the specific innate and adaptive immune responses in each group at challenge, individual genes were examined in their functional groups. In general, the antiviral response was most uniformly upregulated in naïve-vaccinated mice, with naïve-mock vaccinated and preimmune-mock vaccinated mice showing similar levels of antiviral gene expression to each other (**Figure 7**). In response to challenge, all groups showed upregulation of inflammatory cytokines Cxcl10, Cxcl9, and IL-6 and key interferon-stimulated genes (ISGs), such as Oas1a and Oas3, Isg15, Rsad2, Mx1, Ifit2 and Ifit3, and Eif2ak2 (**Figure 7**). However, we observed also differences in certain antiviral response genes. Notably, the IFN-γ-induced chemokine Cxcl10, which signals chemoattraction of T cells, was most highly up-regulated in naïve-vaccinated mice. IFN-γ and IFN-γ induced chemokine Cxcl9 (45, 46) were most strongly up-regulated in preimmune mice, with little differential expression in naïve mice. The inflammasome component Nlrp3 (19, 47) was upregulated in naïve-vaccinated mice, with only minor up-regulation in preimmune-mock vaccinated mice.

**Figure 7.**
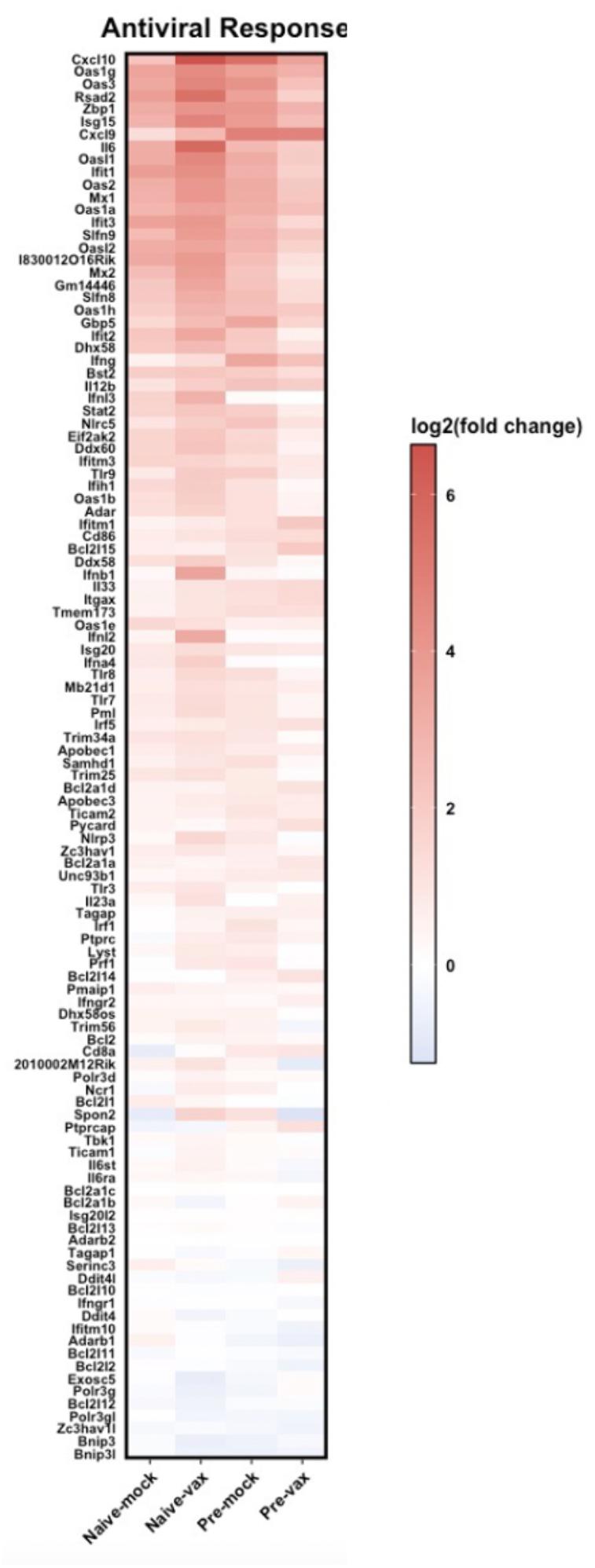
Antiviral immune pathway regulation in mouse lungs at Mex/09 challenge depends on immune background. Heat maps were generated for specific antiviral genes regulated in infected mice per experimental group. The log2(fold change) was plotted for the top differentially regulated genes for gene ontology (GO) biological process (BP) terms “defense response to virus” (GO:0051607), including antiviral innate immune response genes (GO:0140374).

After challenge, all immune backgrounds showed differential expression of genes involved in B-cell mediated immunity (**Figure 8**). For example, B cell chemoattractants Cxcl13 (48, 49) were highly up-regulated in all groups, although minimally in naïve-mock vaccinated mice. B cell genes were most uniformly upregulated in preimmune animals. Both previously infected groups had upregulation of Ighm, encoding the heavy chain of IgM; IL-21, a key regulator of B cell differentiation into plasma cells as well as germinal centre reactions (50); and CD74, which regulates B cell survival (51) (**Figure 8**). Preimmune-vaccinated mice had the greatest up-regulation of IL-13 as well as upregulation of Cd79a/b, a component of the B cell receptor complex (52), suggesting increased BCR signalling which was not observed in the other groups.

**Figure 8.**
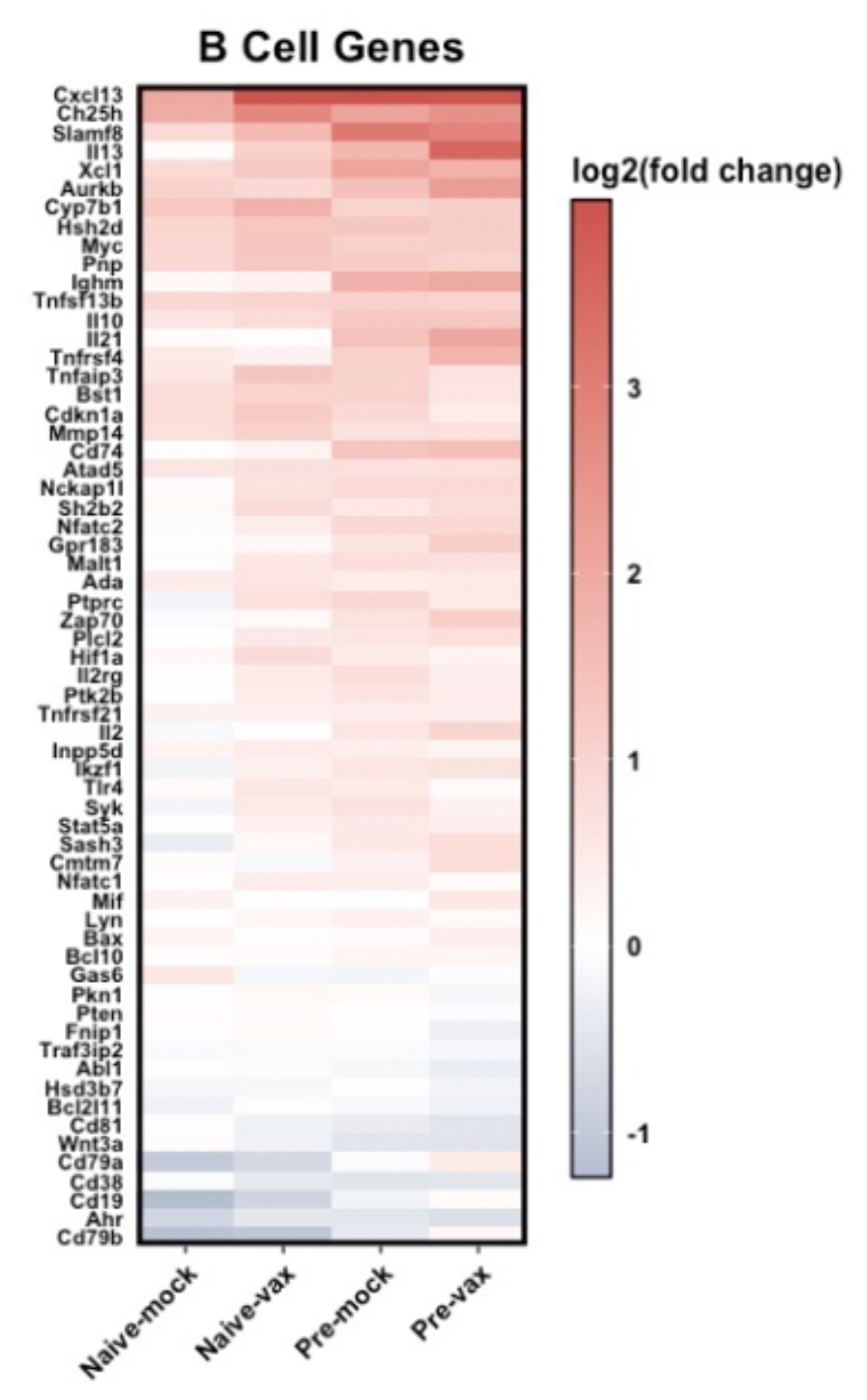
B cell-mediated immune pathway regulation in mouse lungs at Mex/09 challenge depends on immune background. Heatmaps were generated for genes falling under several B cell-mediated immune response pathways. The gene list was generated by combining genes for GO BP terms “B cell chemotaxis” (GO:0035754), “B cell homeostasis”(GO:0001782), “B cell differentiation” (GO:0030183), “negative regulation of B cell differentiation” (GO:0045578), “B cell receptor complex” (GO:0019815), “negative regulation of B cell receptor signalling” (GO:0050859), “B cell apoptotic process” (GO:0001783), and “positive regulation of B cell proliferation” (GO:0030890).

The analysis of genes involved in T cell-mediated cytotoxicity and activation (**Figure 9A**) indicated that all groups had up-regulation of genes involved in T cell cytotoxicity. For example, IL-12b, a subunit of IL-12, as well as granzyme B (Gzmb) were highly upregulated in all groups. All groups also had up-regulation the MHC class I genes H2-M2 and β-2 microglobulin (B2m) (53) where the greatest fold-change occurred in preimmune-vaccinated mice. Key differences were also noted among groups with respect to genes involved in activated T cell proliferation which was most uniformly upregulated in preimmune mice (**Figure 9B**). Strong upregulation of Ido1, which plays a role in suppressing T and NK cells (54, 55) and generating regulatory T cells (56, 57) was identified in preimmune-mock vaccinated mice. Both previously infected groups had upregulation of Pdcd1lg2, encoding programmed cell death 1 ligand 2 which is a key costimulatory molecule for T cell proliferation (58). Il27ra, which contributes to CD4+ T cell differentiation to Th1 cells (59), was also upregulated in previously infected mice but down-regulated in naïve-mock vaccinated mice (**Figure 9B**). Cd24a, which is highly expressed in mouse activated T cells (60). The results of this enrichment and immune pathway-specific gene expression analysis indicated specific immune responses signatures corresponding to each of the four immune backgrounds which we summarized in **Figure 10** in respect to viral protein mutations identified.

**Figure 9.**
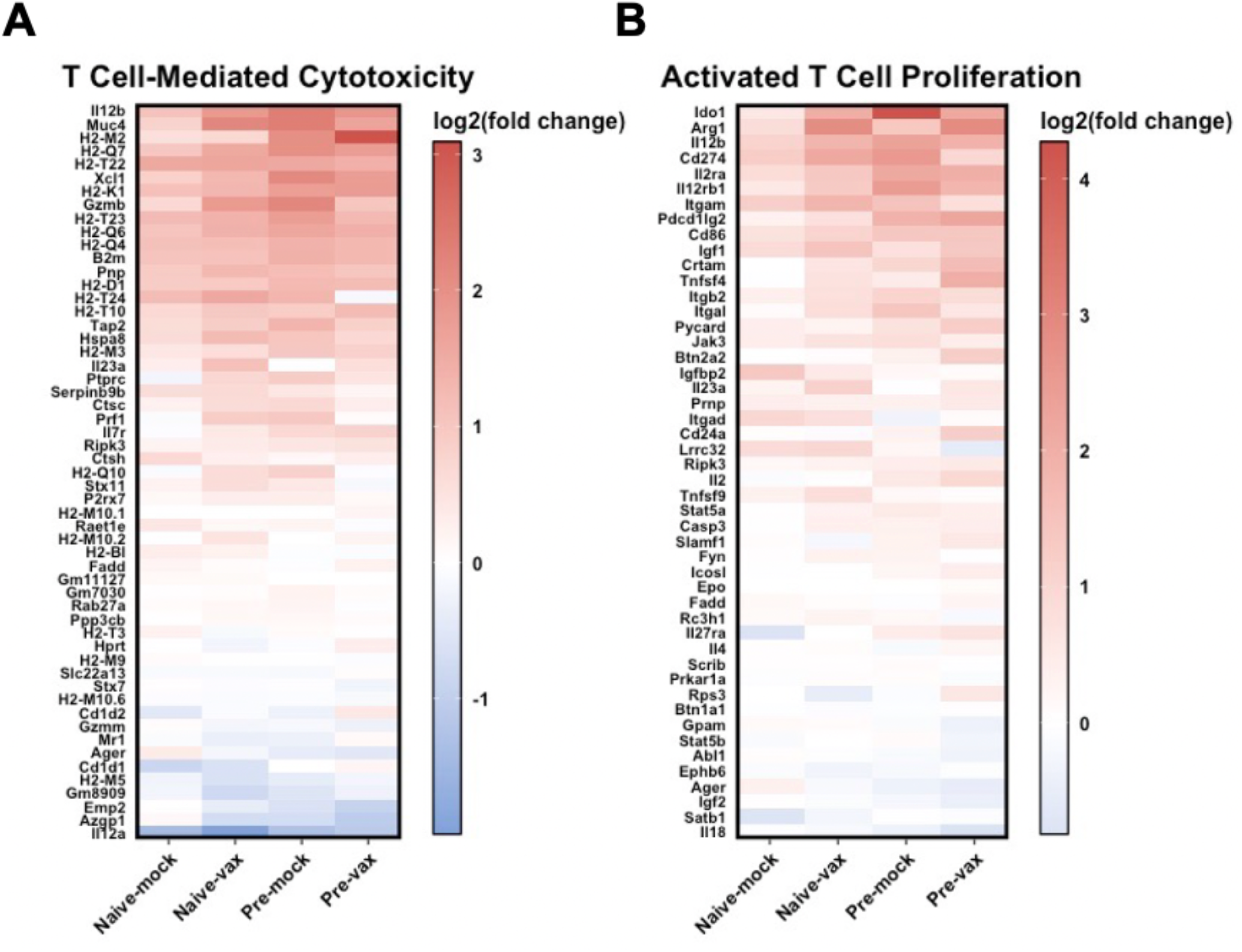
Preimmune-mock vaccinated animals have highest differential expression of genes regulating T cell-mediated cytotoxicity and activated T cell proliferation at three days pc. Heat maps for T cell specific pathways were generated using gene expression data quantified as fragments per kilobases mapped (fpkm), and the log2(fold change) was calculated relative to non-infected mice. The log2(fold change) was plotted for gene ontology (GO) biological process terms “T cell-mediated cytotoxicity” (GO:0001913) and “activated t cell proliferation” (GO:0050798). The heatmaps were generated with ggplot2 package version 3.3.2.

**Figure 10.**
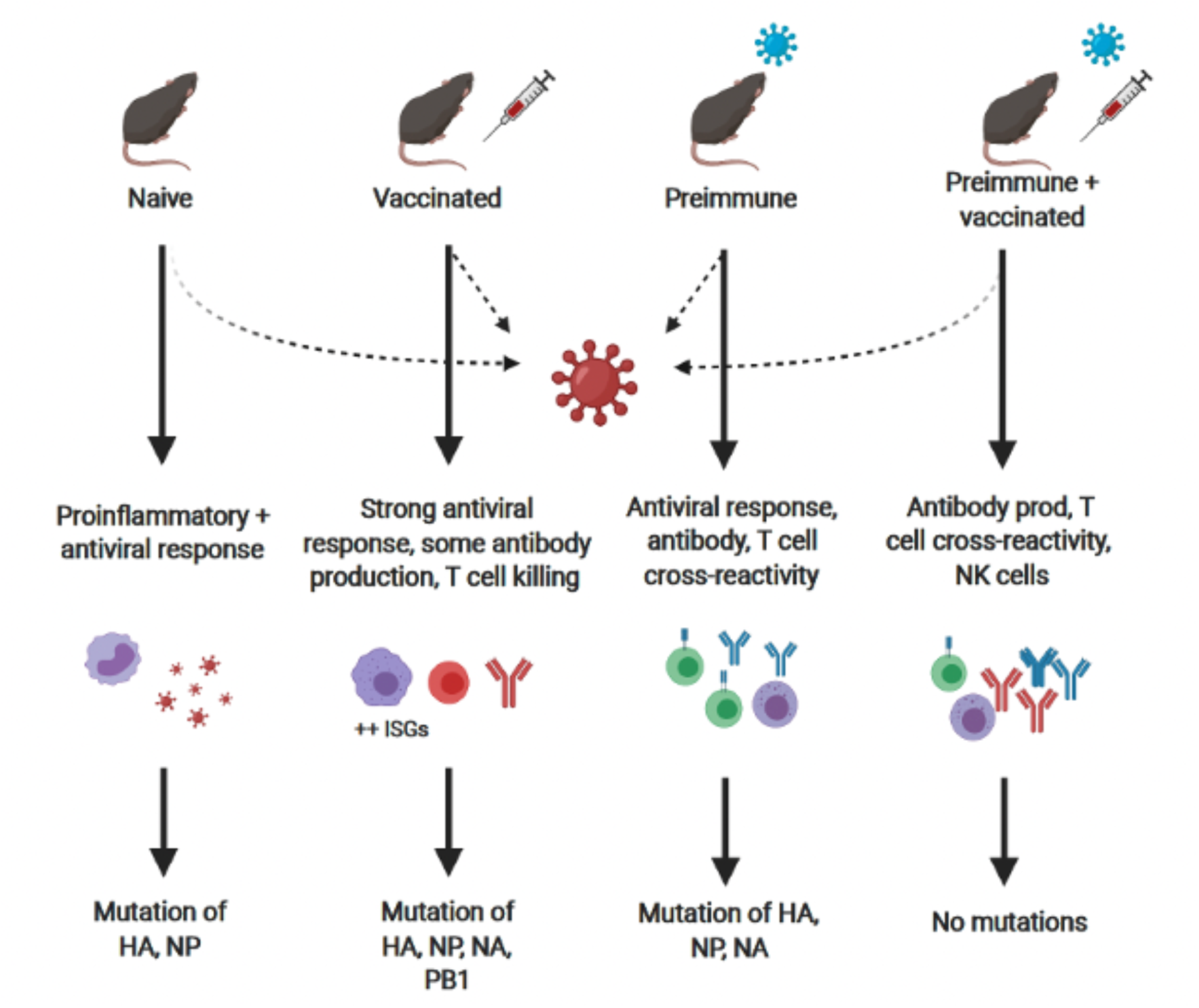
Previous influenza virus infection and vaccination generates selection pressure for influenza virus mutation at challenge. Naïve-mock vaccinated, naïve-vaccinated, preimmune-mock vaccinated, and preimmune-vaccinated animals were challenged with a lethal dose of Mex/09 and lung RNA was extracted at three days post-challenge for viral and host transcriptome sequencing. Signatures of mutations were identified in viral proteins HA, NP, NA, and PB1 that were specific to immune background where each immune background had unique immune responses. At challenge, naïve-mock vaccinated mice had high activation of proinflammatory innate immune mechanisms, which was associated with several mutations on HA (external) and NP (internal). Naïve-vaccinated mice had a significant antiviral response, greater involvement T cell cytotoxicity, and some B cell activation; however, this response facilitates the greatest number of mutations on external proteins HA and NA, as well as internal NP and PB1. Previously infected but not vaccinated mice have the greatest activation of adaptive immune mechanisms which were skewed toward T cell activation and proliferation which was also associated with a higher frequency of viral mutations. Combined preimmunity and vaccination lead to robust antibody production and T cell responses and corresponded to no shared mutations across viral proteins.

## DISCUSSION

The antigenic changes of viral proteins caused by genetic drift is a significant challenge to maintaining protective immunity toward influenza viruses acquired by infection or vaccination. With the goal of understanding the host’s role in antigenic drift, we developed differentially infected and vaccinated mouse models to reflect the diverse immune backgrounds in the human population: no immune history (naive-mock vaccinated), previously vaccinated (naive-vaccinated), previously H1N1 infected (preimmune-mock vaccinated), and previously H1N1 infected and vaccinated (preimmune-vaccinated). When mice of all immune backgrounds were challenged with a contemporary 2009 H1N1 virus that was of a different lineage than the H1N1 virus used for immune background establishment, the RNA was extracted from lung tissue for the analysis of the viral genome as well as the host transcriptome. Our results consistently demonstrated that the largest number of viral mutations as well as the most impactful mutations on T cell and B cell epitopes were from hosts that had only a single viral antigen exposure; that being through a single vaccination or a single previous infection. Additionally, although the single exposed hosts were associated with the greatest viral mutations, the transcriptome analysis indicated unique immune response signatures from each host suggesting different immune pressures to be elicited on the virus during challenge. Previous vaccination in the naive-vaccinated animals led to increased B and T cell activation gene signatures at challenge, which correlated to the greatest number of mutations in the viral genome across all viral proteins analyzed (HA, NA, NP, and PB1). Previously infected mice (vaccinated or mock-vaccinated) had the highest activation of adaptive immune mechanisms at challenge which were skewed toward T cell responses and T cell proliferation along with some genes associated with antibody production. Importantly, the mice with combined previous infection and vaccination had no significant mutations across viral proteins while completely naive infected animals did not survive challenge and had the greatest lung viral titres but also had moderate amounts of viral mutations associated with innate immune responses. The results of our study highlight the important link between the infection and vaccination history of the host and the ever-changing antigenicity of influenza viruses. These results may be informative for next-generation vaccine design as well as anticipatory vaccine strategies.

The use of next generation sequencing (NGS) strategies for influenza viral genomes has allowed scientists to gain a better understanding of the mechanisms driving genome mutations as well as the changes in frequency of viral mutations over time. Influenza viruses have both external and internal proteins that differentially stimulate or suppress the host immune response (reviewed in (61). The immunodominant HA protein accumulates the most mutations across antigenic sites however it is the primary target for host neutralizing antibodies generated at vaccination because of its role in viral entry (14, 62). In one study, mutations in HA including K123N, D131E, K133T, G134S, K157N, and G158E were associated with viral escape due to antibody pressure suggesting the need to understand the mechanisms driving mutations occurring due to antibody pressure (63). NA is also highly immunogenic and can be targeted by antibodies (64). Given their immunodominant nature and propensity to mutate, we hypothesized that HA and NA would have the greatest degree of viral mutation in our study. In agreement with our hypothesis, we found the greatest number of non-synonymous mutations to be in viral genomes extracted from vaccinated only mice as well as the second highest frequency of mutations in mice that had been previously infected but not vaccinated. Interestingly, these mutations were not only abundant on the HA and NA genes as we hypothesized but were also found on the NP and PB1 genes. Specifically, the naïve-vaccinated animals had mutations concentrated in the RBD of HA including W150, L151P, and S183P (65, 66). Another mutation on the RBD mutation, P182Q, was found in the preimmune-mock vaccinated animals only, indicating a mutational pattern distinct from naïve-vaccinated animals. Non-RBD mutations may also be important in conserved and other functional domains of HA (67-70). Two mutations were specific to naïve vaccinated animals (K43N and Y351), and only one in preimmune-mock vaccinated animals (G350V). In a previous study in humans, similar trends of increased viral mutations occurring in vaccinated individuals was observed. H3N2 patient viral isolates were sequenced and identified significant increases in amino acid changes across the HA1 subunit in infected-previously vaccinated patients compared to infected-unvaccinated patients possibly suggesting immune pressure differentially affected the balance of viral populations and/or mutations during viral replication (71). The importance of strain-specific adaptive immunity was more evident when we directly compared HA mutations from naïve-vaccinated mice to those found in preimmune-mock vaccinated mice. Preimmune-mock vaccinated mice, which have antibodies specific to the FM/47 seasonal H1N1 virus as well as cross-reactive T cell immunity had fewer RBD-specific mutations than naïve-vaccinated animals.

We also investigated other immunologically relevant proteins in the influenza virion. Compared to HA, NA showed less mutation in all groups. C292S in the active site of the NA head was identified in all experimental groups, except for preimmune-vaccinated mice. Interestingly, mutations on the stock of the NA protein were also detected but only in animals that did not receive a vaccination (E47G and Q51E in naïve-mock vaccinated; I38T in preimmune-mock vaccinated), however why this would be is unclear. Previous infection led to a high number of mutations in the challenge viral RNA but these were restricted to the more conserved influenza virus proteins NP and PB1. As NP induces strongly cross-reactive CD4+ and CD8+ T cells (37, 39, 72), it is possible that heterologous H1N1 challenge could elicit NP-specific immune memory leading to immune selection of mutations. The greatest degree of immune background-specific mutations in the NP gene occurred in naïve-vaccinated animals where R31G and R441 however, these mutations have not been previously reported to our knowledge and it is unclear why these would have occurred. Looking to PB1, we found only naïve-vaccinated mice to have non-synonymous mutations. These included L95P mutation in the cRNA promoter binding site (73), as well as D617A, G622R, and L624F in the vRNA promoter binding site (44), and S712Y at a core interaction site with PB2 (74). Together, these findings suggests that prior vaccination, but not previous infection, influenced the presence/frequency of mutations on PB1. It is possible that since mutations were found in cRNA and vRNA promoter binding sites, these mutations may provide a fitness advantage when HA is targeted after vaccination.

The impact of pre-existing immunity can be best understood by comparing the transcriptional responses at challenge of our animals with various pre-existing immune backgrounds which included combinations of previous infections and vaccinations. At challenge with Mex/09, the naïve-mock vaccinated animals upregulated ISGs including Cxcl10, Oas1, Irf1, Rsad2, and proinflammatory cytokines IL-6, IL-8, and IFN-g, a signature previously shown in naive ferrets infected with Mex/09 suggesting the classical activation of antiviral responses (75). Moreover, the preimmune-mock vaccinated animals upregulated IgM and Cd8a suggesting early induction of B and T cell-mediated immune mechanisms, which was also found in a ferret influenza reinfection study indicating a recall of cross-reactive previously acquired adaptive immune memory (75). When comparing infection to vaccination, it is known that influenza subunit vaccinations induce more humoral activation and antibody elicitation with little specific cytotoxic T cell immunity (76). Interestingly, our transcriptome analysis at three days post-H1N1 challenge found naïve-vaccinated mice had robust activation of humoral immunity at challenge indicating that B cell immunity was preferentially recalled at live infection as well. Although vaccination significantly decreased viral load after challenge in the lungs compared to mock-vaccinated animals, the vaccinated group had the highest number of mutations across the viral genes analyzed. Conversely, at Mex/09 challenge, preimmune-mock vaccinated animals had early activation of antiviral defense mechanisms, followed by strong upregulation of genes associated with specific T cell activation and B cell proliferation, which are thought to be protective possibly due to resident memory T cells in the lungs (39, 77) and directed toward conserved influenza virus proteins that are targets of cross-reactive T cell responses, such as NP as previously shown (38, 72). Despite similar viral titer, preimmune-mock vaccinated and preimmune-vaccinated mice showed differences in immune response activation at lethal H1N1 challenge. As expected, preimmune-vaccinated mice had the greatest activation of B cell-mediated immune response pathways including B cell receptor signaling and B cell proliferation, as well as activated T cell proliferation. This combined pre-exiting immunity of infection and vaccination was most effective at preventing escape mutations as well as replication advantageous mutations in HA, NA, NP, and PB1. This finding suggests the importance of engaging T cell immunity as a means of removing viral variants as well as specific and non-specific humoral responses to limit new variants. Conversely, it was interesting that the naive-mock vaccinated animals had a low frequency of viral mutations despite highest viral load was immunologically characterized by high proinflammatory immune responses and negligible adaptive responses.

Despite the development of the first influenza virus vaccine many decades ago, influenza viruses still cause significant morbidity and mortality in the human population. Antigenic drift remains the greatest challenge facing influenza virus vaccine development, and although it is generally known that viral mutation is driven by several factors, the exact mechanisms surrounding viral evolution remain largely unexplored in the context of host immunity driving viral mutation selection. Our study provides a comprehensive analysis of specific host infection and vaccination history in relation to influenza viral mutation. Our findings suggested that a having a single form of pre-existing immunity, either from vaccination or infection, favored changes in the frequency of viral mutations that may lead to new viral variants. This study informs an important aspect of host-pathogen interaction that is not considered in traditional vaccine design and may be important especially in the context of other antigenically divergent respiratory viruses such as SARS-CoV-2 and related variants.

## MATERIALS AND METHODS

### Ethics statement

All animal work was completed in accordance with the Canadian Council of Animal Care guidelines. Animal use protocols were approved by the Animal Care Committee for the Dalhousie University Carleton Animal Care Facility (Halifax, NS, Canada; animal ethics number 18-091). Procedures were performed under short-term 3% isoflurane anesthesia to minimize distress. Animals removed for sample collection or humane endpoint reasons were euthanized under 5% isoflurane.

### Experimental animals

Female C57BL/6J mice (5-8 weeks old) were obtained from Jackson Laboratories (Bar Harbor, MN, USA) and housed in HEPA-filtered cage racks adherent to CL2 animal guidelines (Dalhousie University Carleton Animal Care Facility, Halifax, NS, Canada) under standard conditions. Following viral infection, vaccination, and challenge, mice were monitored daily for weight, survival, and other clinical signs of illness. In accordance with the Animal Care Committee, mice were humanely euthanized if 80% of baseline weight was reached.

### Viruses and experimental timeline

All virus work was conducted in a CL2 animal facility. For use in preimmune inoculations, historical mouse adapted H1N1 strain A/Fort Monmouth/1/1947 (FM/47) as well as A/Mexico/4108/2009 (Mex/09) were obtained from the American Type Culture Collection (ATCC) through Cedarlane (Burlington, ON, Canada). The Sanofi FLUZONE quadrivalent influenza vaccine (QIV) for the 2018-2019 season was used for vaccinations (Sanofi Canada; North York, ON, Canada) containing H1N1 (A/Michigan/45/2015 X-275), H3N2 (A/Brisbane/1/2018 X-311), influenza B Yamagata lineage (B/Phuket/3073/2013), and the influenza B Victoria lineage (B/Colorado/6/2017-like virus). Mice were inoculated intranasally with FM/47 at 10^3.5^ EID_50_ (50 uL total) and or vaccinated 50 uL of the human dose of Sanofi QIV in the hind caudal muscle. On day 105, mice were challenged intranasally with Mex/09 10^6^ EID_50_.

### Antibody and viral load assessment

Hemagglutinin inhibition (HAI) titers in mouse serum were determined by HAI assay (28). Antisera were treated for 4 h with Receptor Destroying Enzyme at 37oC and serially diluted in PBS in a 96-well V-bottom plate with 8 hemagglutinin (HA) units/50 uL of antigen. Turkey red blood cells were washed and diluted to 0.05% (vol/vol) in PBS and 50 uL were added to each well, incubated for 30 m, and assessed for agglutination. The HAI titer was calculated as log base 2 of the highest serum dilution factor required to prevent agglutination. Live viral titer in the lungs was measured by TCID_50_ followed by hemagglutination (HA) assay (78). The viral load was calculated according to the Reed and Muench method (79).

### RNA extraction and virus whole-genome sequencing

Total RNA was extracted from the lungs of influenza virus-infected mice using the RNeasy Mini Kit (QIAGEN, Cat. No 74004) according to manufacturer’s instructions. Viral RNA from viral stocks were extracted using QIAamp Viral RNA Mini Kit (QIAGEN, Cat. No. 52906). Influenza viral genome segments were reverse-transcribed using the iScript™ Reverse Transcription Supermix for RTqPCR (Bio-Rad, Cat. No. 1708840) and the Uni12 primer (5’-AGC AAA AGC AGG-3’) (80) for 5 m at 25oC, 20 m at 46oC, and 1 m at 95oC. Viral complementary DNA (cDNA) was then synthesized by PCR for 40 cycles using Taq polymerase (New England Biolabs, Cat. No. M0273) and primers containing Illumina Nextera Transposase adaptors: R1-Uni12 (5’-TCG TCG GCA GCG TCA GAT GTG TAT AAG AGA CAG AGC GAA AGC AGG-5’) and R2-Uni13 (5’-GTC TCG TGG GCT CGG AGA TGT GTA TAA GAG ACA GAG TAG AAA CAA GG-3’) (adaptor and barcode oligonucleotide sequences from Illumina, Inc., San Diego, CA, USA). Annealing and extension steps were performed for 30 s at 55oC and 7 m at 72oC, respectively and PCR products were cleaned using the QIAquick PCR Purification Kit (QIAGEN, Cat. No. 28104). Libraries were sequenced by CGEB-IMR (Centre for Genomics and Evolutionary Biology Integrated Microbiome Resource, Halifax, NS, Canada) (http://cgeb-imr.ca)) in a 300 + 300 bp paired-end MiSeq run (Illumina 600-cycle v3 kit, Cat. No. MS-102-3003). Sequences are available at the Sequence Read Archive (SRA); BioProject ID PRJNA787976; https://www.ncbi.nlm.nih.gov/biosample?Db=biosample&DbFrom=bioproject&Cmd=Link&Link Name=bioproject_biosample&LinkReadableName=BioSample&ordinalpos=1&IdsFromResult=7 87976.

### Viral sequence SNP analysis

Paired-end reads of variable length were imported into Geneious Version 2019.2.1 (Biomatters LTD) (30). Reads were filtered and trimmed according to default settings, and primer sequences were removed. Within Geneious, paired reads were merged using the BBMerge too, and duplicate reads were removed using the DEDupe tool according to default settings. The remaining reads were then mapped to reference sequences for A/Mexico/4108/2009 viral segments. The reference sequences used for each segment alignment are as follows: PB2 (GQ379815), PB1 (GQ149652), PA (GQ149653), HA (GQ223112), NP (GQ149655), NA (GQ149650), M2 (GQ149657), and NEP (GQ149658). Following alignment, single nucleotide polymorphisms (SNPs) above 1% frequency with a minimum of 5 reads supporting their discovery were detected using the SNPs/variants tool. Group-specific SNPs were analyzed using MEGAX (Molecular Evolutionary Genetics Analysis, Version 0.1) (32) was used in intensive modelling mode (33). Structural images of viral proteins were created with PyMOL (The PyMOL Molecular Graphics System, Version 2.3.5, Schrödinger, LLC) and the web-based software Missense3D (34) was used to predict the structural impact of each amino acid substitution. A cavity volume expansion or contraction of ≥ 70 Å3 was defined as structural damage to differentiate from minor expansions or contractions.

### Prediction of B and T cell epitopes

Predicted B cell epitopes across influenza virus proteins and SARS-CoV-2 spike variants were assessed using the DiscoTope 2.0 server (http://www.cbs.dtu.dk/services/DiscoTope/_)(35). The amino acid number and DiscoTope score were used to map predicted B cell epitopes in RStudio (version 3.6.1) using the ggplot2 package (version 3.3.2) (81). For T cell epitopes, the likelihood of influenza virus peptide sequences from our study being presented on mouse was predicted using the Immune Epitope Database (IEDB) Analysis Resource NetMHCpan EL 4.1 (http://tools.immuneepitope.org/main/) restricted to the two the MHC alleles H-2-Kb and H-2-Db. To determine immune background-specific epitopes in influenza protein sequences, T cell epitopes (defined as unique MHC allele*peptide combinations) were compared in RStudio using the tidyverse package (version 1.3.0)(82).

### Host transcriptome sequencing and analysis

Total mouse lung RNA was extracted at days 0 and 3 post-challenge using the RNeasy Mini Kit as described above (QIAGEN, Cat. No 74004) and purified with the QIAquick PCR Purification Kit (QIAGEN, Cat. No. 28104). Extracted RNA was sent to Novogene (Sacramento, CA, USA) for sequencing. Gene expression was quantified in terms of fragments per kilobase of transcript per million mapped (FPKM) and fold-change per gene was calculated relative to uninfected mice, and significance determined using analysis of variance (ANOVA) adjusted p-value < 0.05. Lists of significantly differentially expressed genes (DEGs) were compared across experimental groups to generate Venn diagrams.

Biological processes enriched at challenge were analyzed using the Database for Annotation, Visualization and Integrated Discovery (DAVID) version 6.8 (https://david.ncifcrf.gov/home.jsp) functional annotation tool. The top Gene Ontology (GO) biological processes were selected and compared across experimental groups. The antiviral response (GO term “defense response to virus” (GO:0051607)), T cell proliferation and cytotoxicity (terms “T cell-mediated cytotoxicity” (GO:0001913) and “activated t cell proliferation” (GO:0050798)), and the B cell response (GO terms “B cell chemotaxis” (GO:0035754), “B cell homeostasis” (GO:0001782), “B cell differentiation” (GO:0030183), “negative regulation of B cell differentiation” (GO:0045578), “B cell receptor complex” (GO:0019815), “negative regulation of B cell receptor signalling” (GO:0050859), “B cell apoptotic process” (GO:0001783), and “positive regulation of B cell proliferation” (GO:0030890)) were selected to build heatmaps in RStudio using the ggplot2 package (version 3.3.2). Differential expression was expressed in terms of log2(fold change).

### Statistical analysis

All statistical analyses were conducted in RStudio version 3.6.1. For normally distributed data, one-way analysis of variance (ANOVA) followed by Tukey’s Honest Significant Difference (HSD) was used to determine differences between groups. A p-value threshold of 0.05 was used. For determination of differentially expressed genes by RNA-sequencing, an adjusted p-value threshold of 0.1 was used. All plots were generated in RStudio using the ggplot2 package version 3.3.2 unless otherwise stated.

## Acknowledgments

We greatly acknowledge the critical review and assistance from Dr. Cynthia Swan.

## Contributions

Conceptualization: AAK; Investigation: AAK, MR, MEF, AY, AG, MM; Analysis: AAK, MR, MEF, AG, MM, AY, AS, BX, JD; Writing: AAK, MR, MR, JD; Funding: AAK

## SUPPLEMENTAL FIGURES

**Figure S1.**
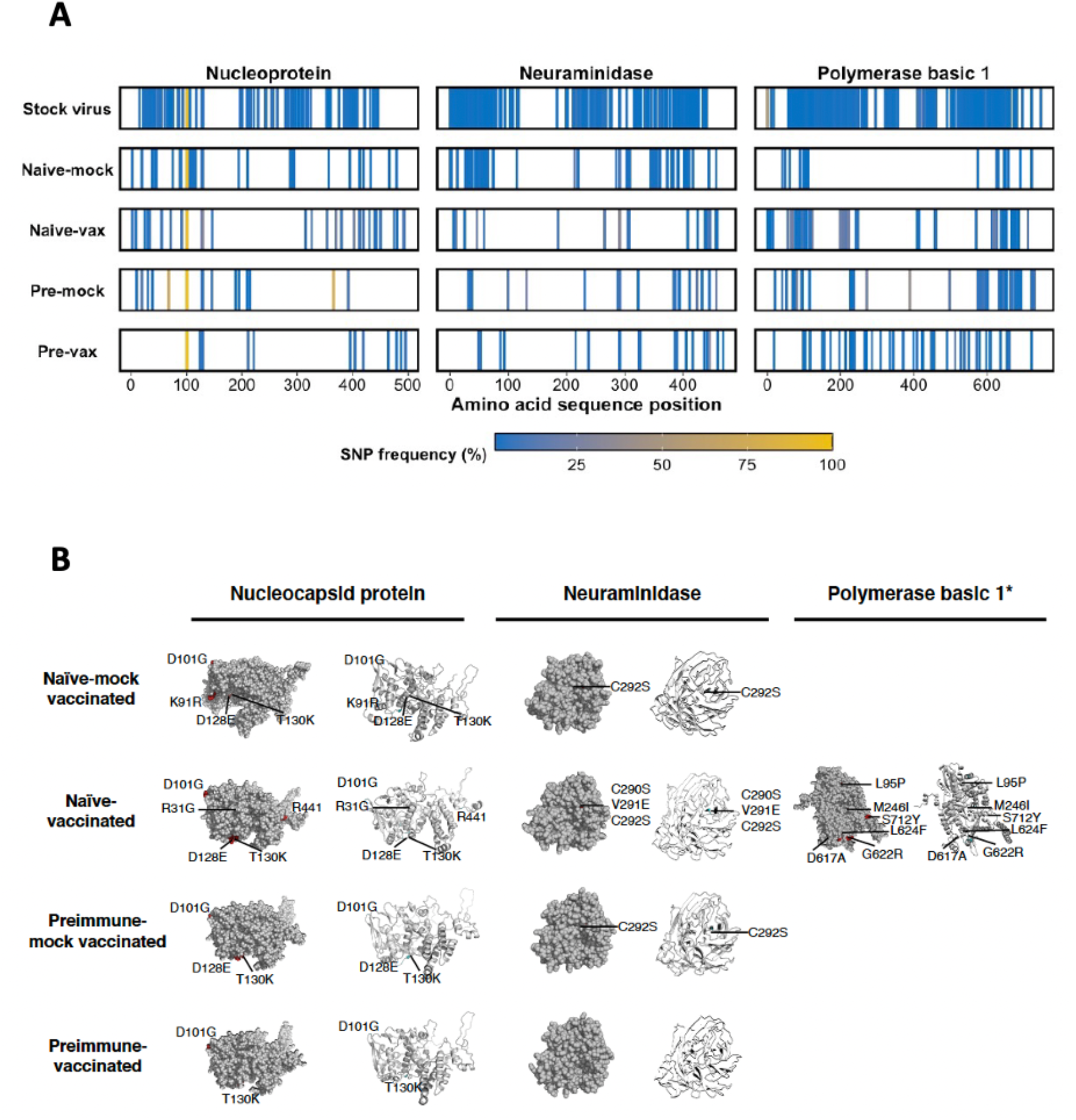
Location and frequency of SNPs on influenza virus nucleoprotein (NP), neuraminidase (NA), and polymerase basic 1 (PB1) amino acid sequences after Mex/09 challenge in mice of different immune backgrounds. (A) Viral RNA sequences were aligned to reference Mex/09 NP, NA, and PB1 using Geneious (NCBI accession numbers GQ149655 (NP), GQ149650 (NA), and GQ149652 (PB1)) and SNPs were identified as done for HA. (B) Structural and antigenic impact of each immune background-specific amino substitution was generated as done previously for HA for each additional protein per immune background. Only amino acid substitutions shared within animals of a group were used to generate folding predictions. *The transmembrane (stalk) domain of neuraminidase was not included in the model.

**Figure S2.**
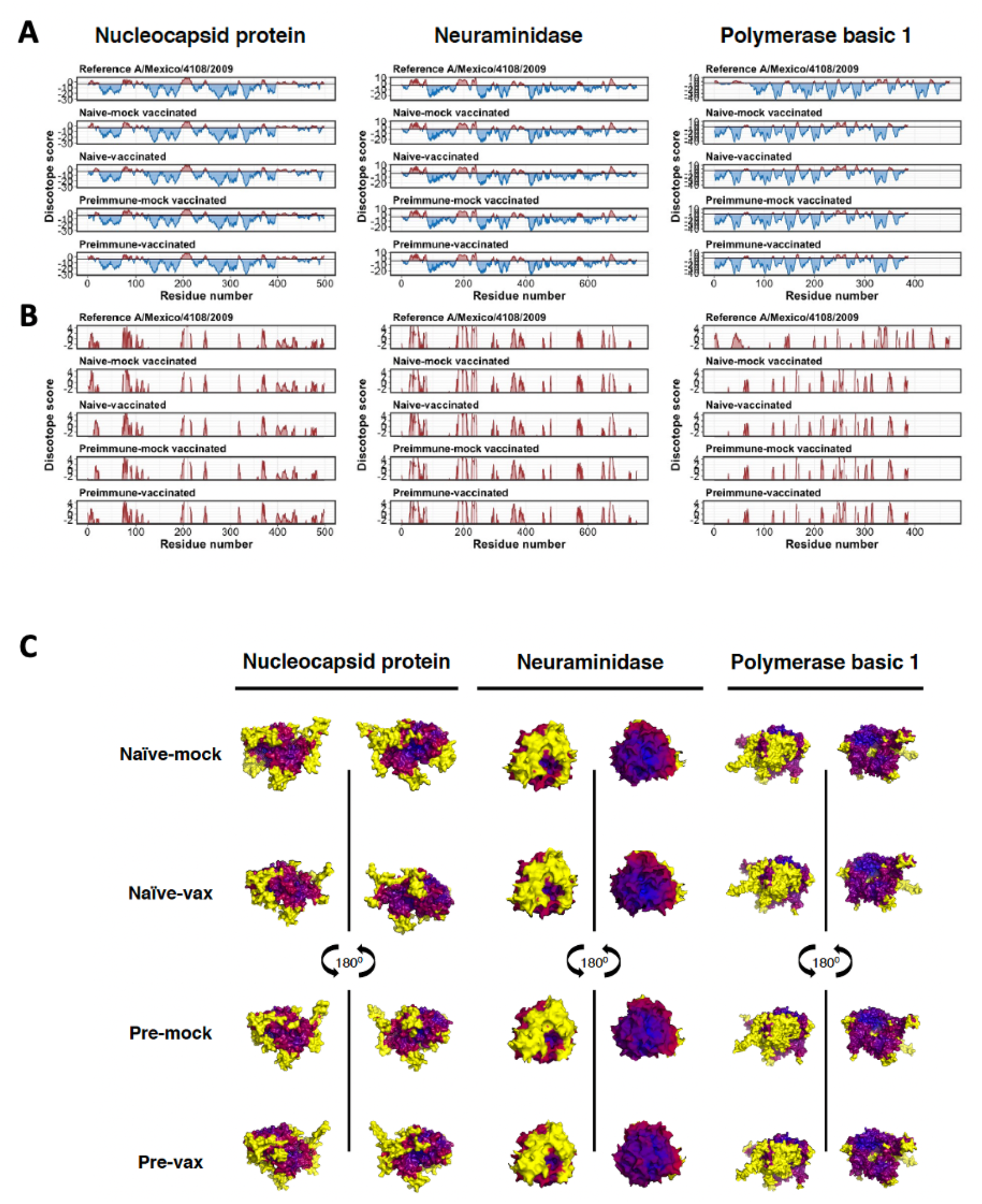
Predicted B cell epitopes on influenza virus nucleoprotein (NP), neuraminidase (NA), and polymerase basic 1 (PB1) after Mex/09 challenge in mice with varying immune backgrounds show minimal differences between groups. Following structural analysis, surface epitopes along NP, NA, and PB1 were predicted using DiscoTope 2.0 as previous. (A) DiscoTope scores falling below (blue) and above (red) the B cell epitope prediction threshold of −3.7 (0.47 sensitivity, 0.75 specificity) were mapped against amino acid sequence number for NP, NA, and PB1 to show regions of likely and unlikely B cell epitopes. Epitope maps for each immune background are compared to that of the respective Mex/09 reference protein, which are as follows: NP (GQ149655), NA (GQ149650), and PB1 (GQ149652). (B) Only the positive B cell epitope prediction results are shown for greater resolution of the B cell epitope regions. (C) DiscoTope scores are shown as heatmaps along the folded protein, with residues colored according to their predicted score as done previously: yellow indicates positively predicted B cell epitopes, red indicates high-scoring residues, and blue indicates low scoring regions.

## SUPPLEMENTAL TABLES

**Table S1.** Summary of immune-background specific mutations detected on influenza viral nucleoprotein (NP), neuraminidase (NA), and polymerase basic 1 (PB1) and predicted structural imapact at three days post-challenge.

| Nucleoprotein (NP) |  |  |  |
| --- | --- | --- | --- |
| Group | SNP | Protein Subdomain | Missense3D prediction |
| Naïve-mock vaccinated | D101G | Body | None; present in stock virus |
|  | K91R | Body | Slightly contracts cavity volume |
|  | D128E | Body | None detected |
|  | T130K | Body | Largely expands cavity volume |
| Naïve-vaccinated | R31G | Body | Slight cavity expansion |
|  | D101G | Body | * |
|  | D128E | Body | None detected |
|  | T130K | Body | Largely expands cavity volume |
|  | R441 | Bottom/tail region | NA |
| Preimmune-mock vaccinated | D101G | Body | * |
|  | D128E | Body | None detected |
|  | T130K | Body | Largely expands cavity volume |
| Preimmune-vaccinated | D101G | Body | * |
|  | T130K | Body | Largely expands cavity volume |
| Neuraminidase (NA) |  |  |  |
| Group | SNP (N1 Numbering) | Protein Subdomain | Missense3D prediction |
| Naïve-mock vaccinated | E47G | Stalk | Contract cavity volume |
|  | Q51E | Stalk | Expands cavity volume |
|  | C292S | Head | Disulfide bond breakage |
|  | S350 | Head | Deletion |
| Naïve-vaccinated | C290S | Head | Disulfide bond breakage and expands cavity volume |
|  | V291E | Head | Introduces buried hydrophilic and charged residue and expands cavity volume |
|  | C292S | Head | Disulfide bond breakage |
| Preimmune-mock vaccinated | I38T | Stalk | Contract cavity volume |
|  | C292S | Head | Disulfide bond breakage |
| Polymerase basic 1 (PB1) |  |  |  |
| Group | SNP | Protein Subdomain | Missense3D prediction |
| Naïve-vaccinated | L95P | cRNA promoter binding site | Buried proline introduced, cavity altered and buried/exposed switch; may affect the cRNA promoter binding |
|  | M246I | Putative nucleotide binding site | No structural damage detected |
|  | D617A | vRNA promoter binding site (C-term) | Salt bridge between ASP 617 and ARG 623 disrupted; may inhibit vRNA promoter binding |
|  | G622R | vRNA promoter binding site (C-term) | Exposed charge introduced; may inhibit vRNA promoter binding |
|  | L624F | vRNA promoter binding site (C-term) | May inhibit vRNA promoter binding |
|  | S712Y | C-term (Core interaction with PB2) | Expand cavity volume by 4.968 Å<sup>3</sup> |

